# Protein-protein interactions in neurodegenerative diseases: a conspiracy theory

**DOI:** 10.1101/2020.02.10.942219

**Authors:** Travis B. Thompson, Pavanjit Chaggar, Ellen Kuhl, Alain Goriely, for the Alzheimer’s Disease Neuroimaging Initiative

**Author notes:** Data used in preparation of this article were obtained from the Alzheimer’s Disease Neuroimaging Initiative (ADNI) database (adni.loni.usc.edu). As such, the investigators within the ADNI contributed to the design and implementation of ADNI and/or provided data but did not participate in analysis or writing of this report. A complete listing of ADNI investigators can be found at: http://adni.loni.usc.edu/wp-content/uploads/how_to_apply/ADNI_Acknowledgement_List.pdf.

## Abstract

Neurodegenerative diseases such as Alzheimer’s or Parkinson’s are associated with the prion-like propagation and aggregation of toxic proteins. A long standing hypothesis that amyloid-beta drives Alzheimer’s disease has proven the subject of contemporary controversy; leading to new research in both the role of tau protein and its interaction with amyloid-beta. Conversely, recent work in mathematical modeling has demonstrated the relevance of nonlinear reaction-diffusion type equations to capture essential features of the disease. Such approaches have been further simplified, to network-based models, and offer researchers a powerful set of computationally tractable tools with which to investigate neurodegenerative disease dynamics.

Here, we propose a novel, coupled network-based model for a two-protein system that includes an enzymatic interaction term alongside a simple model of aggregate transneuronal damage. We apply this theoretical model to test the possible interactions between tau proteins and amyloid-beta and study the resulting coupled behavior between toxic protein clearance and proteopathic phenomenology. Our analysis reveals ways in which amyloid-beta and tau proteins may conspire with each other to enhance the nucleation and propagation of different diseases, thus shedding new light on the importance of protein clearance and protein interaction mechanisms in prion-like models of neurodegenerative disease.

**Author Summary:** In 1906 Dr. Alois Alzheimer delivered a lecture to the Society of Southwest German Psychiatrists. Dr. Alzheimer presented the case of Ms. Auguste Deter; her symptoms would help to define Alzheimer’s disease (AD). Over a century later, with an aging world population, AD is at the fore of global neurodegenerative disease research. Previously, toxic amyloid-beta protein (A*β*) was thought to be the *primary* driver of AD development. Recent research suggests that another protein, tau, plays a fundamental role. Toxic tau protein contributes to cognitive decline and appears to interact with toxic A*β*; research suggests that toxic A*β* may further increase the effects of toxic tau.

Theoretical mathematical models are an important part of neurodegenerative disease research. Such models: enable extensible computational exploration; illuminate emergent behavior; and reduce research costs. We have developed a novel, theoretical mathematical model of two interacting species of proteins within the brain. We analyze the mathematical model and demonstrate a computational implementation in the context of A*β*-tau interaction in the brain. Our model clearly suggests that: the removal rate of toxic protein plays a critical role in AD; and the A*β*-tau ‘conspiracy theory’ is a nuanced, and exciting path forward for Alzheimer’s disease research.

## 1 Introduction

Neurodegenerative diseases such as Alzheimer’s (AD) or Parkinson’s (AD) are associated with the propagation and aggregation of toxic proteins. In the case of AD, it was Alzheimer himself who showed the importance of both amyloid-*β* (A*β*) plaques and tau-protein (*τ* P) neurofibrillary tangles (NFT) in what he called the “disease of forgetfulness” [1, 2]. These two proteins are very different. A*β* forms extracellular aggregates and plaques whereas *τ* P are intracellular proteins involved in the stabilization of axons by cross-linking microtubules that can form large disorganized tangles [3, 4]. Since the early 90’s, when it was first formulated, the “amyloid cascade hypothesis” has dominated the search for cures and treatments [5, 6]. According to this hypothesis, an imbalance between production and clearance of A*β*42 and other A*β* peptides is not only an early indicator of the disease but the causing factor for its initiation, progression, and pathogenesis [7]. However, the repeated failures of large clinical trials focussing on the reduction of A*β* plaques has led many researchers to question the amyloid hypothesis and argue for the possible importance of other mechanisms.

One obvious alternative is that *τ* P plays a more prominent role than the amyloid hypothesis suggests. The *τ* P are usually considered as secondary agents in the disease despite the fact that (1) other *τ* P-related diseases (tauopathies), such as frontotemporal lobar degeneration, are mostly dominated by *τ* P spreading [8]; (2) brain atrophy in AD is directly correlated with large concentrations of NFT [9, 10]; (3) *τ* P distribution determines disease staging [11]; (4) lowering *τ* P levels prevent neuronal loss [12]; (5) *τ* P reduces neural activity and is the main factor associated with cognitive decline [13]. These findings may explain the relative lack of clinical improvements after A*β* suppression and the debate between the relative importance of A*β* proteopathy and *τ* P tauopathy in AD [14]. Furthermore, the similarity in mechanism and progression between prion diseases [15] and classical neurodegenerative diseases led to the formulation of the ‘‘prion-like hypothesis” [16, 17, 18, 19, 20, 21] stating that all these protein-related degenerative diseases are characterized by the progressive spreading and autocatalytic amplification of abnormal proteinaceous assemblies through axonal pathways [22].

Since so many cellular mechanisms are poorly understood *in vivo*, the relative importance of different groups of toxic proteins and their possible interactions have not been established. In particular, both *τ* P and A*β* depend upon and modify the cellular environment [16]. Yet, in recent years a number of studies have linked these two anomalous proteins [23] and raised the possibility that protein-protein interactions in neurodegenerative diseases are a key to understanding both spreading and toxicity [24, 25]. According to Walker, for AD “the amyloid-*β*-*τ* nexus is central to disease-specific diagnosis, prevention and treatment” [14].

Specifically, the following crucial observations have been made in AD: (1) tangles in the cortex rarely occur without A*β* plaques [12]; (2) the presence of A*β* plaques accelerates both the formation of *τ* P aggregates [26] and the interneuronal transfer [27] of *τ* P; (3) the presence of *τ* P induces blood vessel abnormalities [28] and induces neuroinflammation through micro- and astro-glial activation [29]; (4) the presence of A*β* can induce the hyperphosphorylation of *τ* P and the creation of toxic *τ* P seeds by disturbing cell signaling via oxidative stress or through plaque-associated cells (such as microglia) or molecules [30, 31, 32, 33]; (5) A*β* and toxic *τ* P target different cellular components, and doing so amplify each other’s toxic effects [23]; (6) *τ* P mediates A*β* toxicity as a reduction of *τ* P prevents A*β*-induced defects in axonal transport [23]; (7) perhaps more anecdotal, it has also been argued that the lack of clear evidence of dementia in non-human primates, despite the presence of A*β* plaques, could be due to a difference in A*β*-*τ* interactions in these species [34]. From these observation, we extract three crucial modes of interaction:

**M1:** The seeding of new toxic *τ* P is enhanced by the presence of A*β*.

**M2:** The toxicity of A*β* depends on the presence of *τ* P.

**M3:** A*β* and *τ* P enhance each other’s toxicity.

Here, our goal is twofold: first to develop modeling and computational tools to study protein-protein interactions at the brain-organ level and second to test the relative effect of these interactions by direct simulation. Typical approaches for organ-size simulation of dementia progression [35] take the form of either continuous models formulated in terms of anisotropic reaction-diffusion equations [36, 37, 38], or discrete systems on the brain’s connectome network. The discrete approach can be further divided into pure-diffusion linear models [39, 40, 41, 42, 43], probabilistic models [44, 45, 46], or deterministic models [47, 48].

A primary result, of interest to the computational biology community, for the current work will be to show: that non-trivial interactions between A*β* and *τ* P can be realized with relatively simple deterministic models and couplings; and that these interactions can lead to effects with physiological interpretations in neurological disease modeling. Moreover, the mathematical analysis will highlight that clearance mechanisms play a key role in destabilizing the system towards proteopathy. We will therefore select the simplest possible, deterministic, protein kinetic model, including a bulk clearance term, that allows for the expression of both a healthy and toxic regime for a single protein; the heterodimer model [49, 50]. One such system will be defined for A*β*, one for *τ* P and the two heterodimer systems are coupled with a single balanced interaction term. We augment the model by adding an stand-alone, first order equation for damage evolution; this equation expresses the deleterious effects of A*β, τ* P and their interactions. Our general approach, following [48] is to study some of the key properties of this continuous model before discretizing it on a network and solving it numerically on the brain’s connectome graph.

## 2 Model

### 2.1 Continuous model

The simplest possible deterministic aggregation model accounting for the interaction of two protein families, each consisting of a healthy and toxic population, is the heterodimer model [49, 51, 50]. In the heterodimer model: a toxic, misfolded seed protein recruits a healthy protein, induces misfolding, and then fragments; ultimately producing two copies of the misfolded toxic variant. The heterodimer model views these three processes as a single step and expresses this step as an overall mean rate of reaction; such rates can be determined experimentally. Indeed, measuring the mean rates of protein self-aggregation mechanisms, providing best-fit mean aggregation dynamics to deterministic models such as the heterodimer model, is a thriving field of contemporary research [50, 52, 53, 54, 55, 56] and our choice of a deterministic model is inspired by such work. Figure 1 demonstrates the primary molecular mechanism of the heterodimer model; the healthy (blue) protein is approached by the toxic (red) protein and undergoes three separate transitions (small arrows) which are treated as a single transition (long arrow) taking a healthy protein to a misfolded, toxic state.

**Figure 1:**
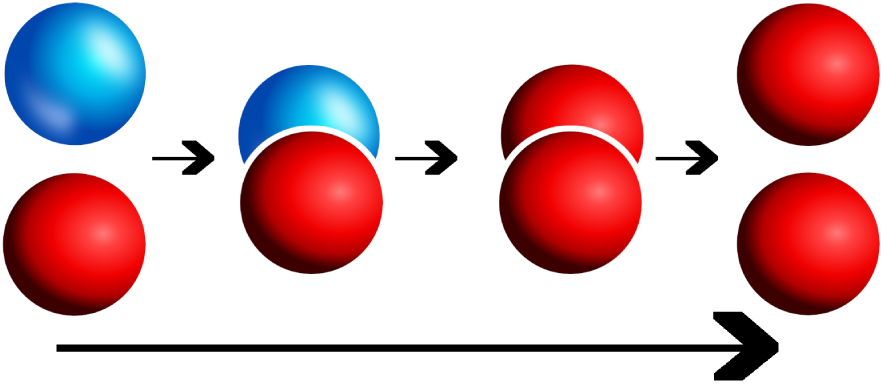
Healthy protein (blue) and misfolded toxic protein (red) transition to two toxic proteins (long arrow) via, from left to right, the kinetics of: recruitment, induced misfolding, and fragmentation.

We are interested in the interaction between two different protein families; motivated by the A*β* and *τ* interactions observed in AD. Towards this end we will consider two heterodimer models: one for A*β* and one for *τ* P. These two models will be coupled together by a single term reflecting that the formation of new toxic *τ* P can be enhanced by the presence of A*β*. The heterodimer model was originally posed [49, 50] as a continuous, non-linear, partial differential reaction-diffusion equation for a single protein. To define the model for our two protein families: let Ω *⊂* ℝ^3^ be a spatial domain of interest and, for **x** ∈ Ω and time *t ∈*ℝ^+^, we denote by *u* = *u*(**x**, *t*), and *v* = *v*(**x**, *t*) the concentration of healthy A*β* and *τ* P. Similarly, we denote by *ũ* = *ũ*(**x**, *t*), and 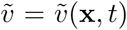, the concentration of toxic A*β* and *τ* P, respectively. Then, the concentration evolution is governed by

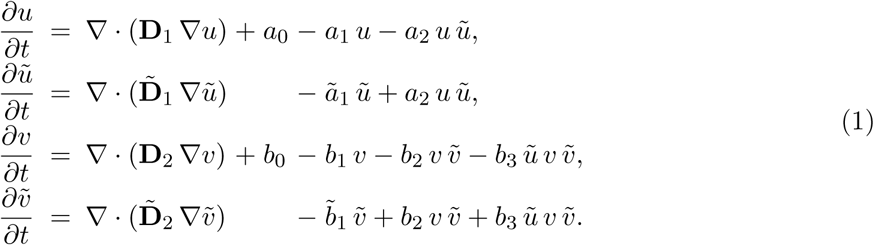

The first two equations, above, correspond to the usual heterodimer model for the healthy and toxic variants of the protein *u*; note that the second heterodimer model, for the variants of protein *v*, deviates from the form of the first by a single balanced term with coefficient *b*_3_. The system (1) could apply to any two families of interacting proteins; though the model is inspired by AD. The parameters are as follows: (*a*_0_, *b*_0_) are the mean production rates of healthy proteins, (*a*_1_, *b*_1_, *ã*_1_,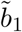) are the mean clearance rates of healthy and toxic proteins, and (*a*_2_, *b*_2_) reflect the mean conversion rates of healthy proteins to toxic proteins. The coupling between the two, otherwise separate, heterodimer models for A*β* and *τ* P, is realized via *b*_3_. The *b*_3_ predicated terms arise from the mode of interaction assumption, c.f. **M1** above, dictating that the presence of A*β* augments the conversion process of healthy *τ* P to toxic *τ* P. We note that toxic A*β* acts as an enzyme in this process and is therefore not depleted. In the absence of production and clearance maps, we assume that all these parameters are constant in space and time. The symmetric diffusion tensors **D**_1,2_ and 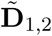 characterize the spreading of each proteins. For isotropic diffusion, these tensors are a multiple of the identity, **D**_1,2_ = *d*_1,2_**1** and ∇· (**D**_1,2_ ·∇(*u*)) = *d*_1,2_Δ(*u*) is the usual Laplacian operator (similarly for *ũ, v* and 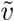) For anisotropic diffusion, the eigenvector with the largest eigenvalue describes the direction of faster diffusion which is used to model preferential propagation along axonal pathways [37].

The coupled system of equations (1) dictates the spread, genesis, and clearance of two healthy species, *u* and *v*, and two toxic species, *ũ* and 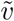, of proteins throughout the domain Ω. The presence of toxic proteins near a point **x** ∈ Ω can disrupt the extracellular environment of neurons near **x** and impair their intracellular function. A broad range of coupled effects can contribute to neuronal impairment; including: chronic inflammation, erosion of the blood-brain barrier surrounding vessels, accelerating tau hyperphosphorylation, disrupting normal synaptic efficacy, and deafferentation, among others. A hallmark of neurodegenerative proteopathies is cognitive decline; propelled by the various coupled effects induced by the presence of toxic aggregates and the widespread erosion of neuronal integrity. The nuanced coupling between these disparate deleterious effects is not well understood; nevertheless, we employ the observation that such effects are generally correlated with larger concentrations of misfolded aggregates to define a gross measure of regional neuronal ‘damage’ denote by *q*(**x**, *t*) ∈ [0, 1]. This damage variable takes the perspective that *q*(**x**, *t*) = 0 signifies that the neurons in a neighborhood of **x** ∈ Ω are functional and healthy whereas *q*(**x**, *t*) = 1 implies that neurons near **x** have reached a fully-degenerate asymptotic state whereby they are either no longer functioning or fully deceased. For the evolution of the damage we assume a simple, first-order rate model:

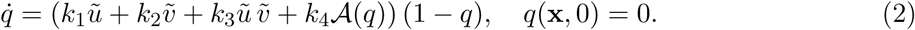

When *k*_4_ = 0: the evolution equation (2) can be seen as a first order reaction model, i.e. exponential decay, for the transformed variable 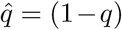; the associated rate of decay is then dependent on the deposition concentration of pattern of *ũ* and 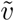. The first two parameters in (2) denote contributions to neuronal dysfunction, near **x**, due to presence of isolated toxic aggregates; while the third term accounts for contributions requiring, or accelerated by, the presence of both toxic species aggregate species together. Thus, the third term engenders both toxic effects M2 and M3; c.f. Section 1. The first three terms of (2) account for neuronal dysfunction in a neighborhood of the point **x** while the last term, 𝒜(*q*), incorporates non-local contributions, such as transneuronal degeneration, whereby the impairment, or death, of neighboring neurons can increase [57] the probability of impairment near **x**; thus leading to an increased mean rate of local decline. This nonlocal term does not have a simple representation within the continuous framework as the positions of neuronal bodies is not explicitly encoded. However, we will see that in the discrete case, there is a natural way to take this effect into account and we will delay the discussion of this term to Section 2.2.

### 2.2 Network model

A simple coarse-grain model of the continuous system can be obtained by building a network from brain data. The construction is obtained by defining nodes of the network to be regions of interest in the domain Ω, typically associated with well-known areas from a brain atlas. The edges of this network represent axonal bundles in white-matter tracts. The brain connectome is then modeled as a weighted graph 𝒢with *V* nodes and *E* edges obtained from diffusion tensor imaging and tractography. A network approximation of the diffusion terms, having the general form ∇· (**D** ∇*u*) or similar, in the system (1) will be constructed by means of a weighted graph Laplacian. The weights of the weighted adjacency matrix **W**, used to construct the graph Laplacian, are selected as the ratio of mean fiber number *n*_*ij*_ by mean length squared, 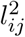, between node *i* and node *j*. That is:

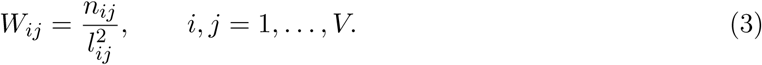

The choice of weights, above, are consistent with the inverse length-squared dependence incurred by canonical discretizations of the continuous Laplace (diffusion) operator appearing in (1). The weighted degree matrix is the diagonal matrix with elements

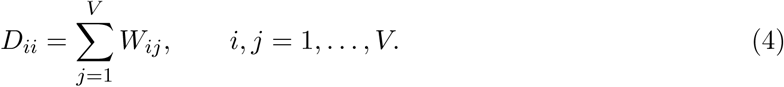

Additionally, we define the graph Laplacian **L** as

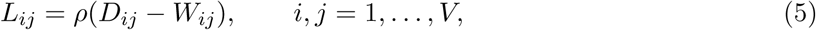

where *ρ* is an overall effective diffusion constant. The adjacency matrix for the simulation is derived from the tractography of diffusion tensor magnetic resonance images corresponding to 418 healthy subjects of the Human Connectome Project [58] given by Budapest Reference Connectome v3.0 [59]. The graph contains *V* =1015 nodes and *E* =70,892 edges and is shown in Figure 2.

**Figure 2:**
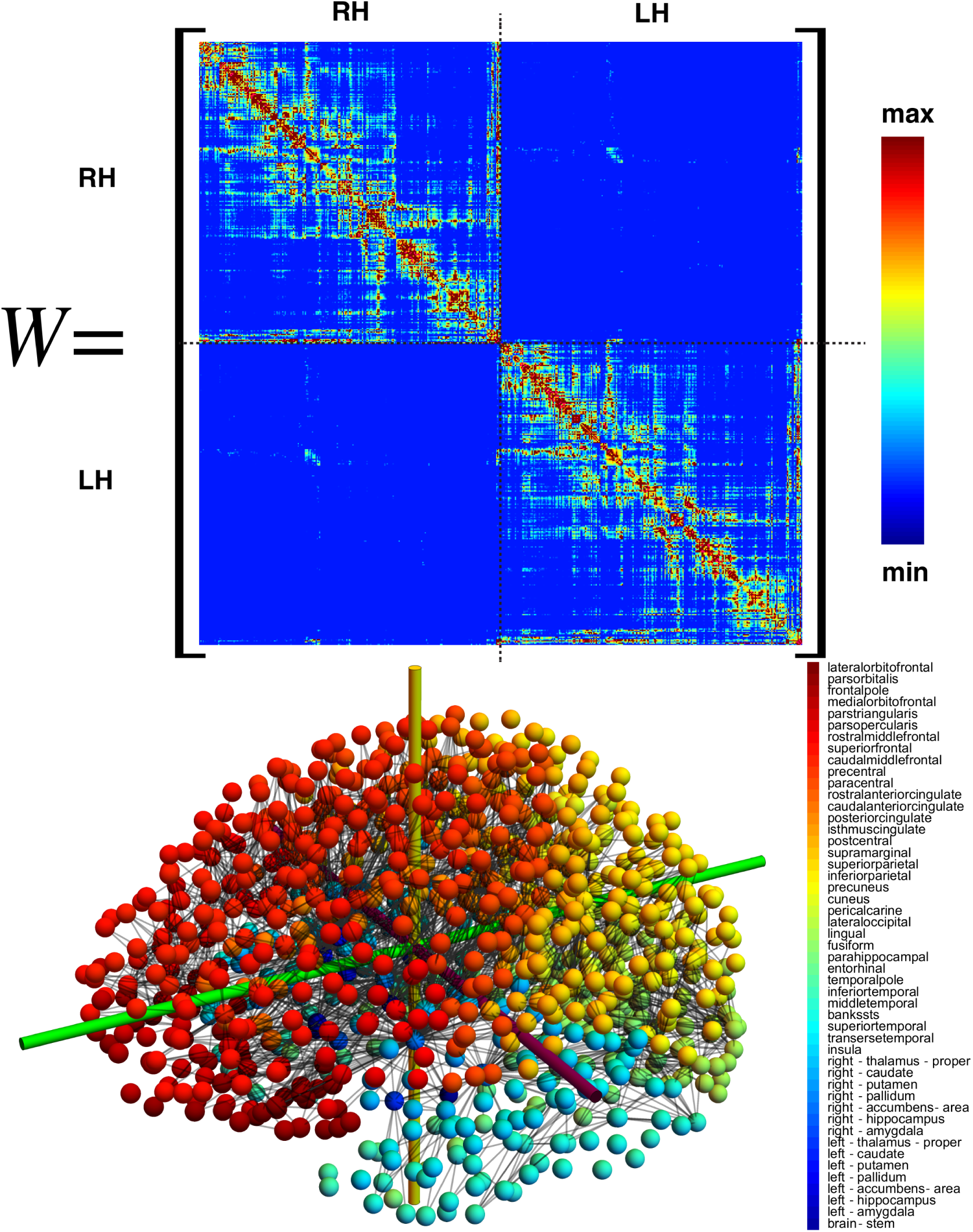
(Bottom left) The average of 419 brain connectomes with *V* = 1, 015 vertices spanning (bottom right) 49 associated brain regions; the strongest 2,773 edge connections are shown. The weighted adjacency matrix (top) corresponding to the averaged connectome (bottom).

Let (*u*_*j*_, *ũ*_*j*_) be the concentration of healthy and toxic A*β* and 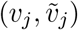 denote the concentration of healthy and toxic *τ* P at node *j*. The network equations corresponding to the continuous model then take the form of a system of first-order ordinary differential equations. There are four such equations, (*u*_*j*_, *ũ*_*j*_, *v*_*j*_, 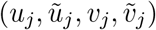), for each of the 1,015 vertices in the system; these four nodal equations are:

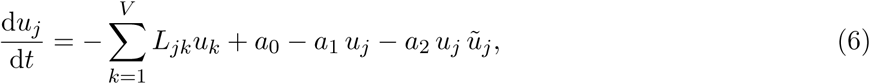

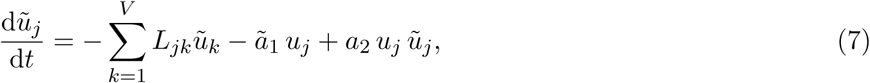

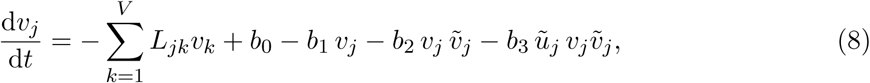

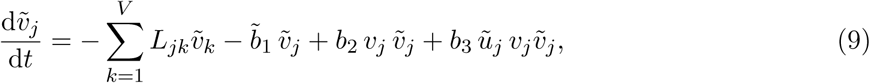

where *j* = 1, *…, V* = 1, 015. Similarly, for the damage model we define a damage variable *q*_*j*_ at each node *j* and assume the same law

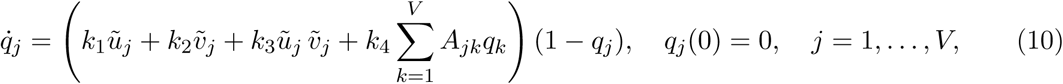

where *A*_*jk*_ is the weighted network adjacency matrix *A*_*jk*_ = *n*_*jk*_*/l*_*jk*_ if *j ≠ k* (and *n*_*jk*_ *>* 0) and 0 otherwise. Thus *k*_4_ has the interpretation of a ‘transmission speed’; the time it takes for the effects of degeneracy in cell *k* to reach cell *j*. The weighting chosen in the adjacency matrix term is inspired by the propagation of transneuronal degeneration from a node to its neighbors.

## 3 Analysis of the continuous model

### 3.1 Homogeneous system

It is instructive to start with an analysis of the homogeneous system obtained by assuming that there is no spatial dependence. This analysis applies to both network and continuous models. In this case, both systems reduce to the dynamical system

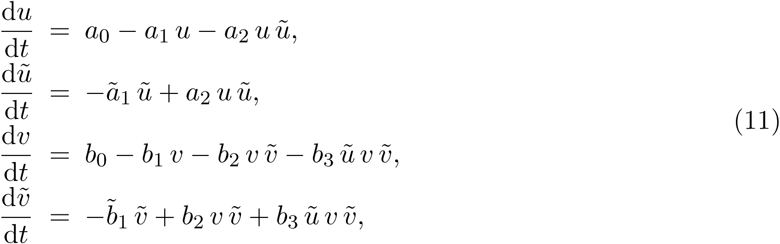

where all variables and initial conditions are assumed to be positive and all parameters are strictly positive.

**Damage evolution** For the homogeneous system above the concentrations remain homogeneous for all time. Damage, in contrast, is node-dependent and expressed by the (nodal) variable *q*_*j*_ ∈ [0, 1]. Indeed, in this case, the non-local term associated with transneuronal degeneration, commensurate with the tensor *A*_*jk*_ in Eq. (10), cannot be homogeneous. Nevertheless, the damage dynamics are simple enough to describe. Damage will initially increases linearly in time, homogeneously, from the initial value *q*_*j*_ = 0. The increase will then trend exponentially at each node, with node-dependent time scales depending on the local node’s degree, and saturate to the value *q*_*j*_ = 1 asymptotically in time at each node.

### 3.2 Stationary points

The stationary points and stability of the homogeneous system (11) are instructive; they inform the disease dynamics implied by the local model. The system (11) can exhibit one, two, three, or four stationary points depending on the parameters; these are:

1. Healthy *τ* P-healthy A*β*: This stationary state is always a solution to (11) and is descriptive of an individual with zero toxic load; no amyloid plaques or neurofibrillary tau tangles. The state is given by:

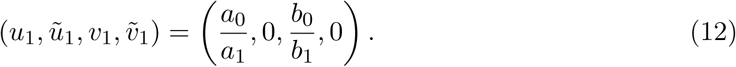
2. Healthy *τ* P–toxic A*β*: This state describes a diseased brain wherein some A*β* plaques exist but the tau fibril (NFT) concentration or that of hyperphosphorylated tau is non-existent or negligible. A description of this stationary state in terms of the base problem parameters is:

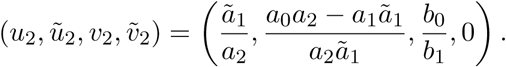

In terms of *u*_1_ = *a*_0_*/a*_1_, from (12), and *u*_2_ = *ã*_1_*/a*_2_ it is given by

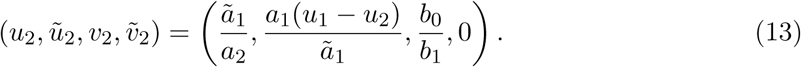

Since the concentrations must be non-negative: the form of *ũ*_2_, above, implies that *u*_1_ *≥ u*_2_. This results in the condition of *ã*_1_*/a*_2_ *≤ a*_0_*/a*_1_. In other words either the clearance term of toxic A*β* must be sufficiently small, the conversion term must be sufficiently large, or a ratio of the two, to allow for the existence of a toxic state.
3. Toxic *τ* P–healthy A*β*: This stationary state is a conceptual dual to the previous state above; granted, toxic *τ* P does not influence the A*β* population whereas A*β* does induce additional *τ* P formation. As in (13) we express this state, immediately here, in terms of *u*_1_ = *a*_0_*/a*_1_ and *v*_1_ = *b*_0_*/b*_1_ as

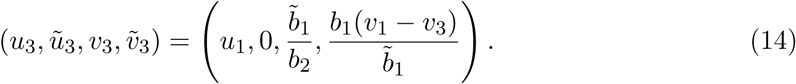

Requiring *v*_1_ *≥ v*_3_ implies that 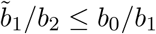.
4. Toxic *τ* P–toxic A*β*: This stationary state reflects the invasion of a patient’s brain by both toxic amyloid beta and toxic tau. As in (12)-(14) we write the state in terms of the previous state variables *u*_1_ = *a*_0_*/a*_1_, *u*_2_ = *ã*_1_*/a*_2_, *ũ*_2_ = *a*_1_(*u*_1_ − *u*_2_)*/ã*_1_, 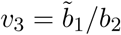 and *v*_4_, defined below, as:

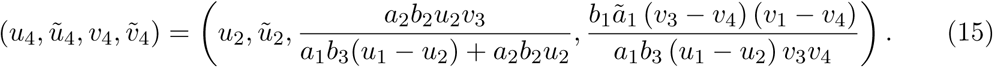

Introducing

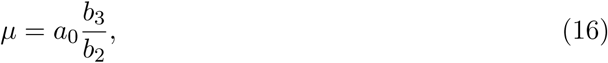

into (15) gives

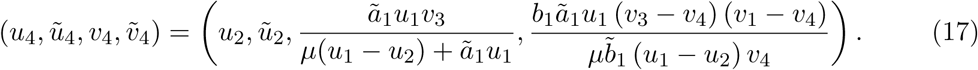

### 3.3 Stability

We briefly discuss the stability of the stationary points. In addition we distinguish between the two possible ‘disease’ phenomena of (11): the case of a disease system characterized by the dynamics of a four-stationary-point model and the case of a disease system characterized by three fixed points.

**Eigenvalues of the linearized system** The linearization of (11) about any fixed point (*u, ũ, v*, 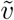) is governed by the Jacobian matrix

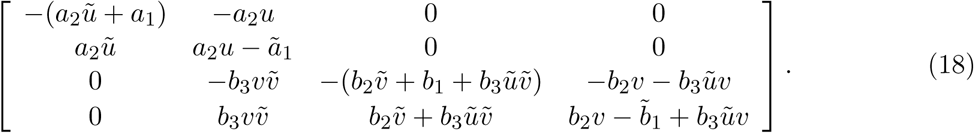

The first two eigenvalues of (18) correspond to the A*β* subsystem, e.g. (*u, ũ*), of (11). Since the coupling of (11) is a one-way coupling these eigenvalues are given by the corresponding eigenvalues of the uncoupled heterodimer model:

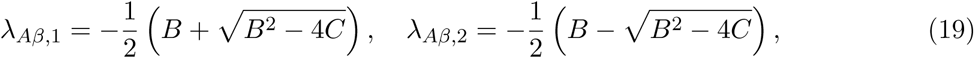

where *B*(*u, ũ, a*_1_, *a*_2_, *u*_2_) = *a*_1_ + *ã*_1_ + *a*_2_ (*ũ* − *u*) and *C*(*u, ũ, a*_1_, *a*_2_, *u*_2_) = *a*_2_ (*ã*_1_*ũ* − *a*_1_*u*) + *ã*_1_*a*_1_. The third and fourth eigenvalues of (18), corresponding to the coupled 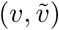 tau system of (11), can be written as

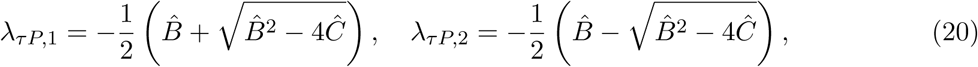

with 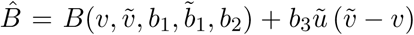 and 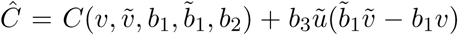. The form of the tau eigenvalues coincides with those for A*β* when *b*_3_ = 0 or when *ũ* vanishes.

### 3.4 Disease phenomenology

We can interpret the different stationary states in terms of disease dynamics and define, accordingly, different disease states.

#### The healthy brain

A healthy patient represents an instantiation of the healthy stationary state whereby 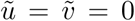. For the Healthy *τ* P-healthy A*β* state to exist we must have *a*_0_ *≤ a*_1_ and *b*_0_ *≤ b*_1_, i.e., (*u, v*) ∈ [0, 1] *×* [0, 1] are valid concentrations. A failure in healthy clearance, either with an amyloid clearance value satisfying 0 *≤ a*_1_ *< a*_0_ or with a tau clearance of 0 *≤ b*_1_ *< b*_0_, implies the non-existence of a physically relevant healthy state (c.f. (12)). It is instructive to note that the expressions *a*_0_*/a*_1_ and *a*_2_*/ã*_1_ (respectively *b*_0_*/b*_1_ and 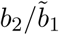) express a balance of healthy A*β* production to clearance and toxic A*β* production to clearance (respectively healthy *τ* P and toxic *τ* P production to clearance). Consider the following balance of clearance inequalities:

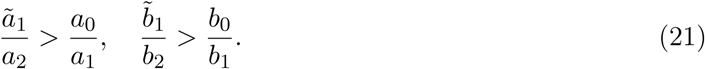

A patient satisfying (21) enjoys full stability to perturbations while in the healthy state (12). That is: if (21) holds with (*u, ũ, v*, 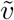) given by (12) then the real parts of the eigenvalues (19)-(20) are negative and the production of small amounts of toxic A*β*, or of toxic tau, or the excess production of healthy A*β*, or healthy tau, results in a quick return to the healthy homeostatic baseline state of (12). The above implies that the model (6)–(9) recognizes the critical role that clearance plays in neurodegenerative diseases. A low value of toxic clearance *ã*_1_, respectfully 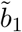, with sustained healthy clearance or a low value of healthy clearance *a*_1_, respectfully *b*_1_, with sustained toxic clearance is enough to trigger an instability capable of driving the system away from the healthy state.

**The susceptible brain** From the previous discussion, we conclude that an unfavorable alteration in clearance mechanisms not only renders the healthy state unstable to perturbations but brings into existence the other stationary points characterizing various pathological conditions.

Indeed, a well established clinical biomarker for Alzheimer’s disease is a drop in soluble amyloid concentration in the cerebrospinal fluid; directly suggesting a decrease in *a*_1_. Recent evidence also suggest [60] that toxic tau filaments in chronic traumatic encephalopathy patients enclose hydrophobic molecules which may contain blood-born pathogens; a possible result of vascular damage from an impact. Such a finding could imply, for instance, that repeated traumatic injury causes vessel rupture and a subsequent proclivity for this unique form of toxic tau production. The stage is then set to trigger a pathological decline when the critical relation (21), corresponding to tau, is violated due to a balance of increased toxic load and age-induced clearance deficit.

The moment of susceptibility occurs when the inequality of (21) becomes an equality. Mathematically, this parameter configuration is a transcritical bifurcation for the homogeneous system (11) at the coincidence of a combination of the states (12)-(14). Clinically, this is the point whereby additional stationary states are physically meaningful and pathology development becomes a possibility.

#### The proteopathic brain

The proteopathic brain has suffered a perturbation from the healthy stationary state; due to the instability in the system this patient is progressing towards a diseased state. The potential pathology phenotypes depend on the patient’s individual parameter values. In particular, if *ã*_0_*/a*_1_ *≥ a*_1_*/a*_2_ holds then the existence of (13) is physically meaningful and if 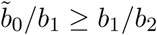 holds then the same is true of (14). It may be the case, depending on the combination of failed clearance subsystems and specific predisposition for toxic loading, that both relations hold simultaneously.

A necessary (clinical) existence criterion for the proteopathic stationary point (15) can be observed directly from the equation for 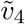 in (17): namely

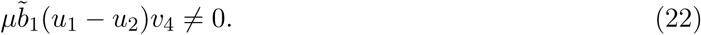

This implies that the parameter *b*_3_, defining *µ* in (16), cannot vanish.

Finally since *b*_3_ ≠ 0 and the numerator of of *v*_4_, in (17), is always non-negative we see that (15) always exists when *u*_1_ *> u*_2_ and when both *v*_3_, *v*_1_ *≥ v*_4_ or when both *v*_3_, *v*_1_ *≤ v*_4_. An important observation is that, though the modeling of the pathology of (15) is tied to that of (13) it is not inextricably tied to (14); this is due to the fact that we may always choose *b*_3_, c.f. (16), such *v*_4_ is smaller than both *v*_3_ and *v*_1_. Thus, with a suitably strong A*β* tau-toxification interaction the state (14) is not needed in order to produce tau proteopathy; that is, the model admits a pathology whereby toxic tau is created solely by the presence of toxic A*β*. Therefore, there are two clinically interesting patient proteopathies for our analysis: the case where the patient model consists of all four disease state equilibria, (12)-(15), and the case where the patient model has the three equilibria (12), (13) and (15).

#### Primary tauopathy

In this case, all four equilibria exist which requires both *ã*_1_*/a*_2_ *< a*_0_*/a*_1_ and 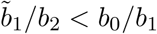. An example of this dynamics is shown in Figure 3. We see that the presence of toxic A*β* always implies a higher level of *τ* P. Indeed, we have

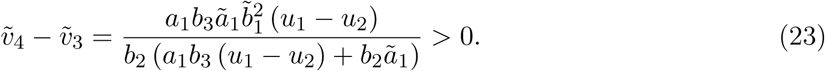

We refer to this case as *primary tauopathy* as the invasion due to *τ* P exists *independently* of A*β*. The effect of A*β* is to increase the concentration of toxic *τ* P and, possibly, increase the associated damage.

**Figure 3:**
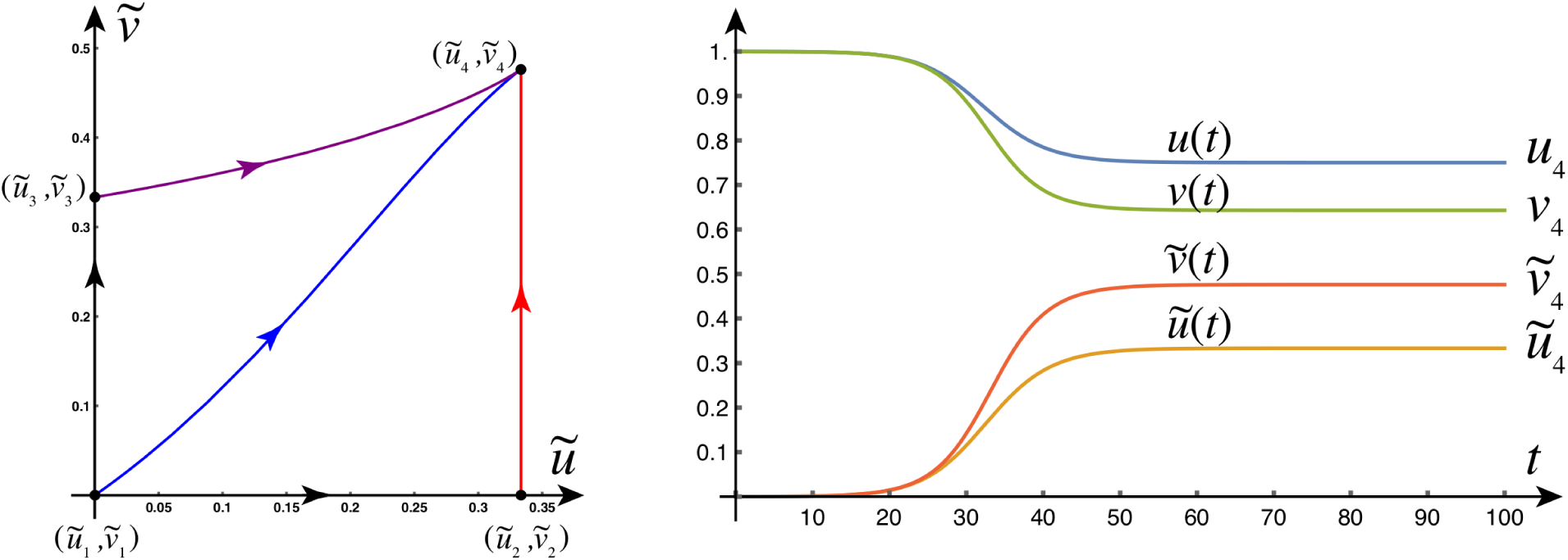
(Left) Phase plane (*ũ*, 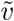) with four equilibria. Homogeneous dynamics of the toxic states. Note that this is a two-dimensional slice of the four-dimensional phase space. (Right) When four different states co-exist, only the fully toxic state is stable as shown by the time-dynamics plot. (Parameters: *a*_0_ = *b*_0_ = *a*_1_ = *a*_2_ = *b*_1_ = *b*_2_ = 1, 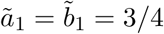, *b*_3_ = 1*/*2).

#### Secondary tauopathy

In secondary tauopathy the evolution of *τ* P depends on the primary invasion of A*β*. Parameters corresponding to secondary tauopathy can be obtained by choosing *ã*_1_*/a*_2_ *< a*_0_*/a*_1_ and 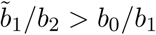 (hence, 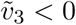) while taking *b*_3_ large enough so that 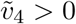.

It is useful to outline key observations, explored further in the manuscript, regarding the nature of the dependence of *τ* P pathology 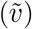 on the presence of A*β* pathology (*ũ*) in this regime. In Section 3.5 we will see that the onset of toxic *τ* P follows from the presence of toxic A*β*. We will also see, in Section 3.5, that the speed of toxic *τ* P propagation appears to be limited by the speed of propagation of toxic A*β*; c.f. Figure 6 for both observations. This is distinctly different than the case of primary tauopathy where toxic *τ* P and toxic A*β* can evolve separately. Indeed, Figure 3 (left) shows a stationary point of the form 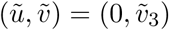 and the fully invaded asymptotic states, Figure 3 (right), satisfy 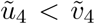; clearly indicating that the additional coupling of *ũ* to 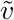, in (9) does not limit tau pathology expression, in primary tauopathy, to that of A*β* pathology.

In light of the apparent dependence of toxic *τ* P spreading (Section 3.5) on toxic A*β* propagation in secondary tauopathy it is instructive to ask: will the asymptotic level of toxic *τ* P pathology concentration be limited by the asymptotic concentration of toxic A*β*? For instance: Figure 4 (right) seems to indicate that this is the case. However, we will see, in Section 5.1.2, that the asymptotic state value of 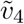 is not limited by that of *ũ*_4_. Figure 16a shows that 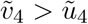 is possible depending on the parameter selection. This observation leads naturally to a further question: is it a strict requirement of secondary tauopathy to have a toxic A*β* concentration at all? That is: can we have 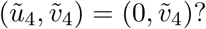 Suppose this is the case: then, according to (12) and (15), we have *ũ*_4_ = *ũ*_2_ = 0 so that *u*_1_ = *u*_2_. When *u*_1_ = *u*_2_ we have, again from (15), that *v*_4_ = *v*_3_; this leaves 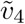, in (15), in an indeterminate form. Thus, to understand 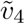 when *u*_4_ = 0, we use that 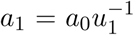, c.f. (12), and consider:

**Figure 4:**
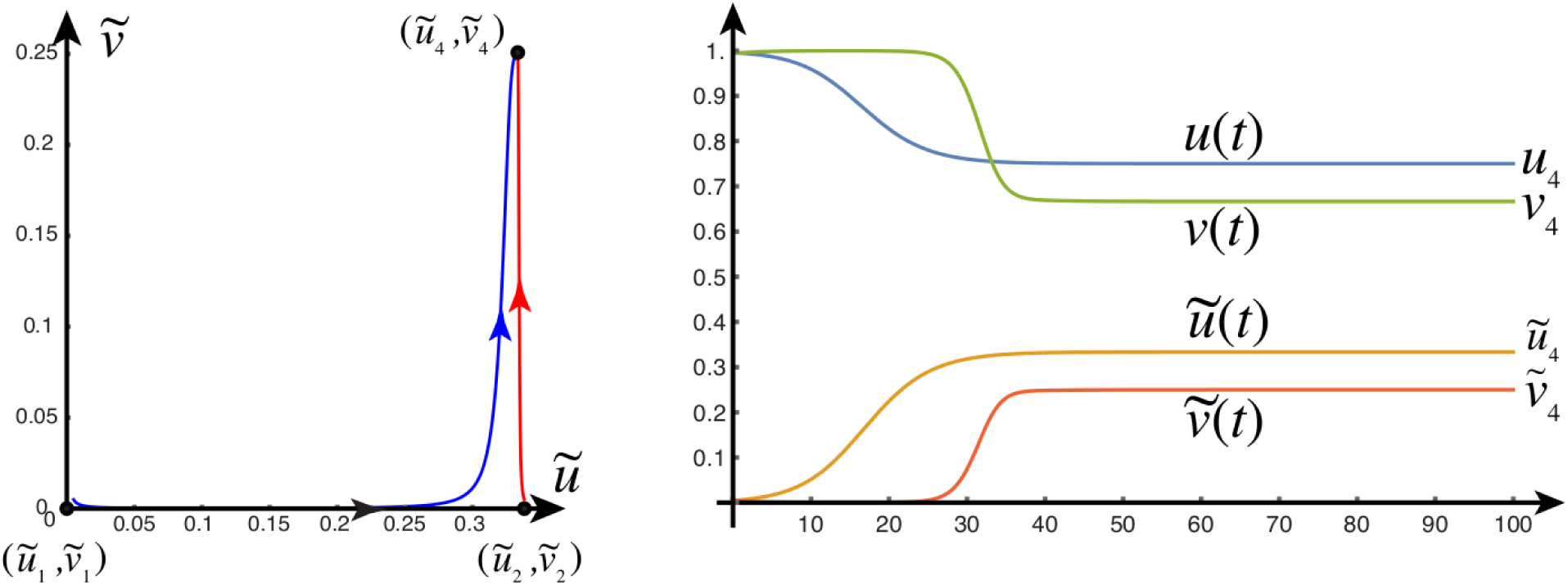
(Left) Phase plane (*ũ*, 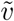) with three equilibria. (Right) When three different states co-exist, only the fully toxic state is stable as shown by the time-dynamics plot. (Parameters: *a*_0_ = *b*_0_ = *a*_1_ = *a*_2_ = *b*_1_ = *b*_2_ = 1, *ã*_1_ = 3*/*4, 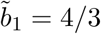, *b*_3_ = 3). Note that trajectories are initialized by taking the initial condition *ϵ* = 0.005 away from an equilibrium point.

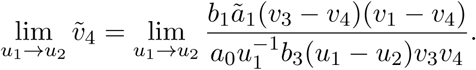

Due to the indeterminate form of the limit above: we apply L’Hôspital’s rule and compute, after simplifying, that

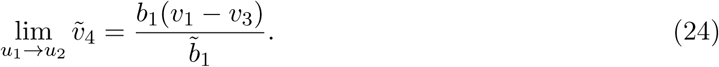

As discussed above: the regime of secondary tauopathy occurs when 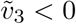; resulting in (14) being an invalid steady state. This directly implies that *v*_1_ − *v*_3_ *<* 0 so that the expression (24) is negative. The above shows that if the asymptotic toxic A*β* concentration, *ũ*_4_, vanishes in the case of secondary tauopathy then the asymptotic toxic tau concentration, 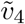 is necessarily negative; this cannot occur in a physical system. To summarize: the sustained presence of a toxic *τ* P population, in secondary tauopathy, *requires* the presence of toxic A*β*; this implies that the computational observations of Section 5.1.2, regarding the reliance of toxic *τ* P development and perpetuation, on the presence of A*β* are strict and not merely one possible method of toxic *τ* P development.

### 3.5 Front propagation

We can explore the spatio-temporal behavior of the system by first considering a reduction to one dimension (Ω = ℝ) and subsequently analyzing the spread of toxic protein via the study of traveling waves. From the theory of nonlinear parabolic partial differential equations, we expect pulled fronts that connect one equilibrium state to a different homogeneous state [49].

First, consider the two uncoupled fronts emanating from the healthy state (*u*_1_, *ũ*_1_, *v*_1_, 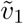) and connecting either to (*u*_2_, *ũ*_2_, *v*_2_, 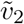) or (*u*_3_, *ũ*_3_, *v*_3_, 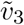). To obtain these fronts, we linearize (1) around the healthy state (*u*_1_, *ũ*_1_, *v*_1_, 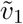) and obtain the decoupled system

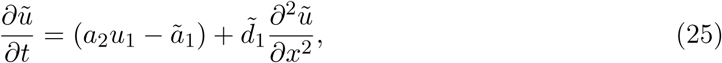

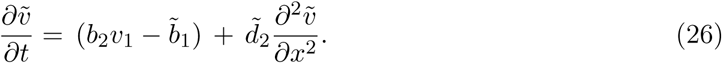

Starting with initial positive data, the system will develop fronts and the asymptotic selected speed is the minimum possible speed for this linear system [61, 62]. Traveling wave solutions to (25)-(26) are obtained explicitly by first performing a traveling wave reduction (*u*(*x, t*) *→ u*(*z*) with *z* = *x* − *ct* and so on for the other variables) and then looking for linear solutions of the form *u* = *C* exp(*λz*) which leads to a family of possible solution with speeds *c* = *c*(*λ*). The smallest such speed is the selected speed for the asymptotic dynamics. In our case, the front speeds are

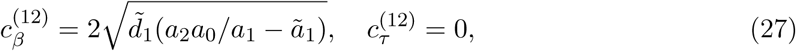

where 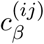 and 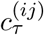 denote the speeds of the front from state *i* to state *j* (whenever such a front exists) for the A*β* fields (*u, ũ*) and *τ* P fields 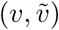, respectively. The front speeds for the second transition are

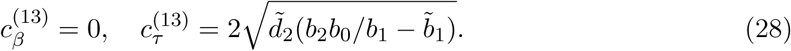

Similarly, if both fields are seeded initially, we have

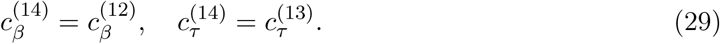

We see that these fronts only exist if *a*_2_*a*_0_ *> ã*_1_*a*_1_ and/or 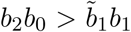 which are the conditions for the existence of toxic states found in the previous section. Trivially, a front between two states can only develop if such states exist.

Second, we consider the possibility of fronts propagating from equilibrium state 2 to state 4. To do so, we linearize the equations around (*u*_2_, *ũ*_2_, *v*_2_, 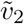) and repeat the previous steps to find

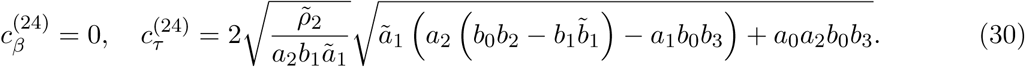

#### Primary tauopathy

As an example of the interactions between the two fronts, we consider a toxic A*β* front on the real axis *x* propagating to the right interacting with a *τ* P front propagating to the left (see Figure 5). They evolve initially with constant speeds 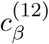 and 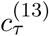 respectively

**Figure 5:**
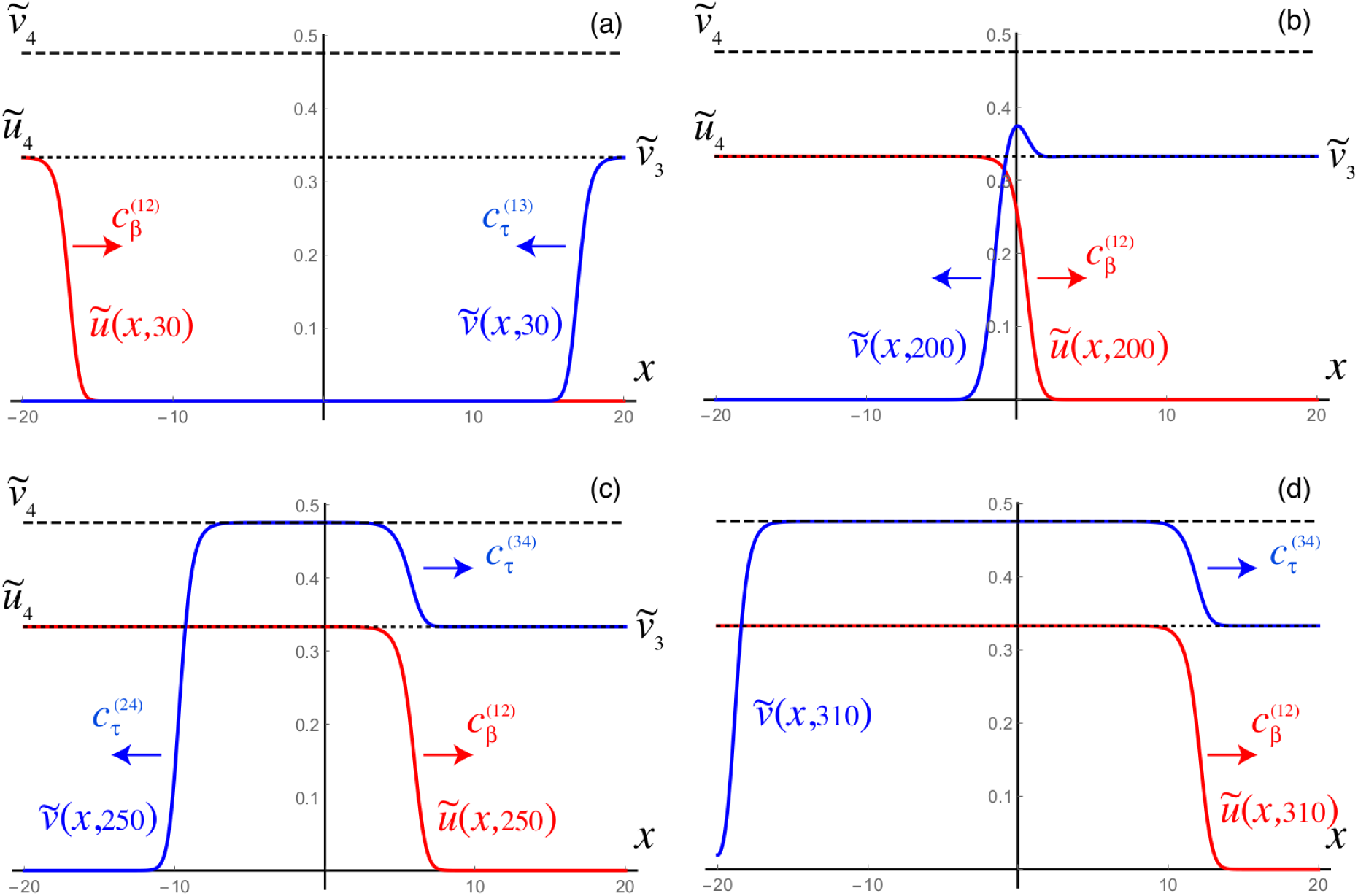
Toxic front dynamics of *ũ*(*x, t*) and 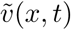 in primary tauopathy. An A*β* front propagating to the right travels towards a *τ* P front propagating to the left (a). The interaction (b) increases the toxic level of *τ* P and creates a second front propagating to the right connecting 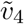 to 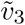 (c). The front profiles are shown at time *t* = 30, 200, 250, 310. Parameters as in Figure 3: *a*_0_ = *b*_0_ = *a* = *a* = *b* = *b* = 1, 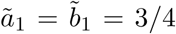, *b*_3_ = 1*/*2, which leads to 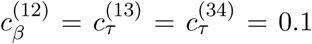 and 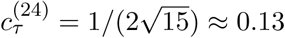. Neumann boundary conditions are used on both sides of the finite interval for all variables.

(Figure 5 top). However, when they overlap, the interaction creates an increase in the concentration of *τ* P (Figure 5 top) which both boosts the front to speed 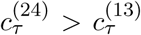 and initiates a new front propagating backward to fill the interval to the global stable equilibrium 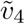 with speed 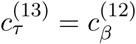. The A*β* front is never affected by the presence of toxic *τ* P.

#### Secondary tauopathy

As a second example, we consider the case where the A*β* front causes the creation of a non-zero toxic *τ* P state (see Figure 6). Initially, a toxic A*β* front propagates to the right in an environment with negligible values of toxic *τ* P (Figure 6a). The passage of the front leads to the rapid expansion of toxic *τ* P (Figure 6b) which evolves at a speed close to 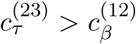 (Figure 6c). Eventually it catches up with the A*β* front where it matches its speed 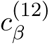 (Figure 6d).

**Figure 6:**
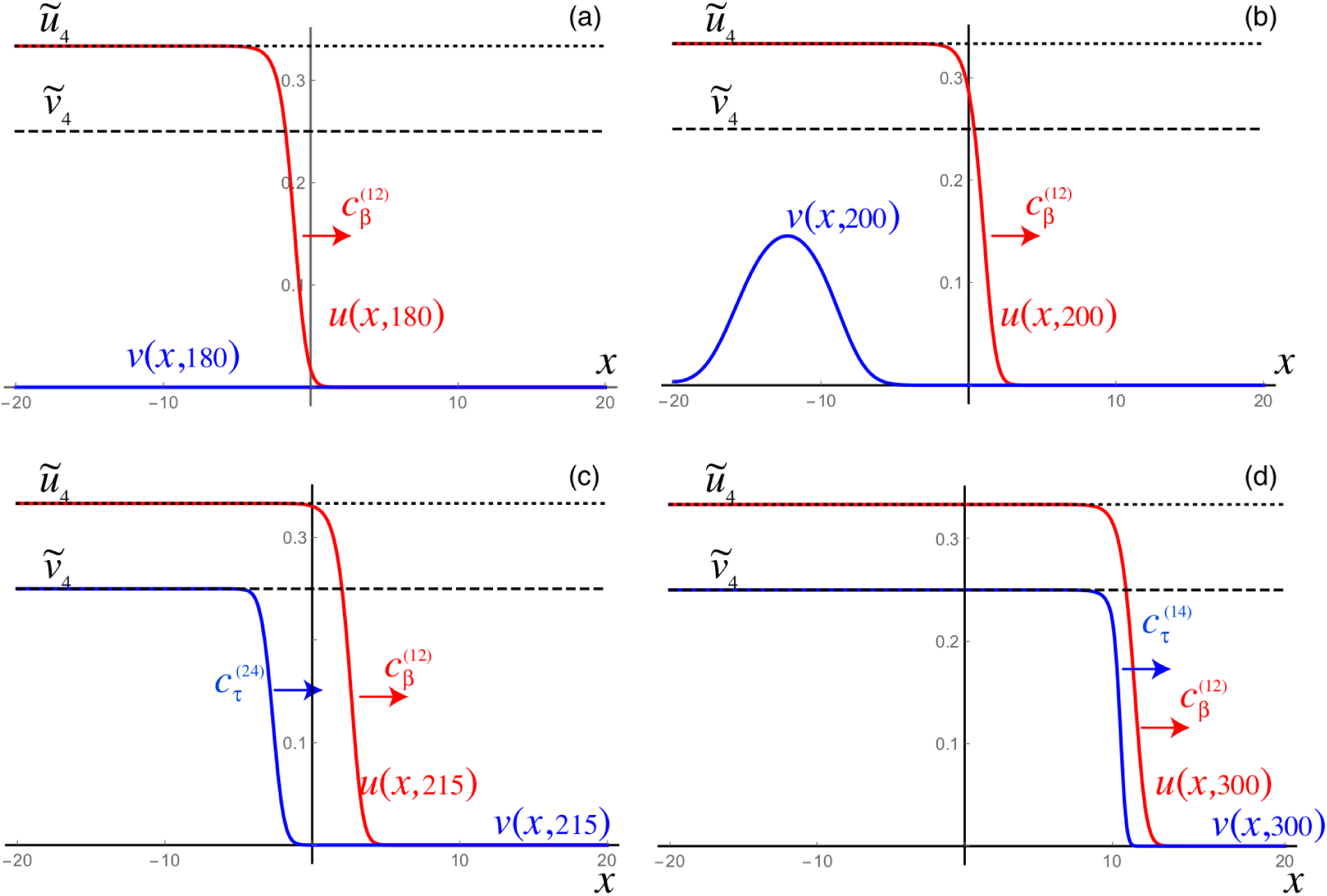
Toxic front dynamics of *ũ*(*x, t*) and 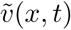 in secondary tauopathy. An A*β* front propagating to the right in a domain with negligible toxic *τ* P (but with a healthy *τ* population). The interaction (b) creates a rapid expansion of the toxic levels of *τ* P and creates a *τ* P front propagating to the left but faster than the A*β* front. Hence, it eventually catches up with the front (d) and matches its speed. The front profiles are shown at time *t* = 180, 200, 215, 300. Parameters as in Figure 4: *a*_0_ = *b*_0_ = *a*_1_ = *a*_2_ = *b*_1_ = *b*_2_ = 1, *ã*_1_ = 3*/*4, 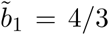, *b*_3_ = 3, which leads to 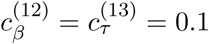 and 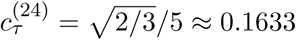. Homogeneous Neumann boundary conditions are used on both sides of the finite interval for all variables. Initial seeding of toxic *τ* P on the positive interval only with 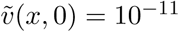 for *x >* 0 and 0 otherwise.

Third, the front propagating from equilibrium state 3 to state 4 is constrained by the evolution of the *u* and *ũ* fields. Therefore, we find

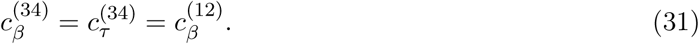

## 4 Network model dynamics

We have established the properties of our system of equations in the homogeneous case and in one-dimension. The study has lead to the identification of two fundamental disease propagation modes depending on the parameters: the primary tauopathy where toxic *τ* P states can exist independently from the A*β* concentration, but are enhanced by its presence; and the secondary tauopathy where the presence of toxic *τ* P is slaved to the existence of toxic A*β*. We can use this analysis as a guide to the simulation of the full network equation. Equations (6)-(10) were discretized on the reference connectome [59], c.f. Section 2.2, using CVODE as part of the SUNDIALS nonlinear ODE solver library [63] in addition to KLU [64] as part of the SuiteSparse [65] linear algebra library. Snapshots of the dynamics are shown in subsequent figures, but full movies can be found in the supplementary material.

As a way to systematically test the validity of our computational platform, we have performed two main tests. First, we reproduce the homogeneous states in the full network and second, we reproduce the transition between homogeneous states. Both tests are detailed in Appendix A in addition to a discussion regarding a choice of hypothetical, non-clinical parameters for illustration purposes; c.f. Appendix A.2 for full details on the parameter selection and the resulting numeric values characterizing each pathology state.

### 4.1 Front dynamics on networks

Propagating front solutions for the system of partial differential equations (1) were considered, via linearization around the healthy state and reduction to one spatial dimension, in Section 3.5. Propagating fronts represent fundamental modes of disease pathology dynamics that can also be realized by the network model of (6)-(10) as we now demonstrate. We consider two different network for front propagation. First, a three-dimensional regular cubic lattice with *n*_*x*_ = 30 nodes in the *x*-direction *n*_*y*_ = 6 nodes in the *y*-direction and *n*_*z*_ = 3 nodes in the *z*-direction, spaced equally at unit length. Second, we use the physiological brain connectome domain of Figure 2, but we choose initial conditions on two sides of the brain to illustrate the front dynamics. In the next section we will consider the same domain but with realistic initial conditions.

#### 4.1.1 Primary tauopathy

The first example is that of primary tauopathy corresponding to the parameters of Table 5.

**Table 1:** Summary of selected ADNI patient group statistics

| | 18F-AV1451 $\tau P$ PET group | 18F-AV45 $A\beta$ PET group |
| --- | --- | --- |
| Total patient count | 38 | 42 |
| Male patients | 25 | 28 |
| Female patients | 13 | 14 |
| Mean age ± SD, years | 78.4 ± 5.14 | 78.4 ± 5.1 |

**Table 2:** Comparison with ADNI Alzheimer’s patient PET data: General synthetic parameters

| Parameter | Value | Parameter | Value | Parameter | Value | Parameter | Value |
| --- | --- | --- | --- | --- | --- | --- | --- |
| Healthy amyloid-β population parameters | | | | | | | |
| $\rho$ | 1.38 | $a_0$ | 1.035 | $a_1$ | 1.38 | $a_2$ | 1.38 |
| Toxic amyloid-β population parameters | | | | | | | |
| $\rho$ | 0.138 | $\tilde{a}_1$ | 0.828 | $a_2$ | 1.38 | | |
| Healthy $\tau P$ population parameters | | | | | | | |
| $\rho$ | 1.38 | $b_0$ | 0.69 | $b_1$ | 1.38 | $b_2$ | 1.035 |
| $b_3$ | 4.14 | | | | | | |
| Toxic $\tau P$ population parameters | | | | | | | |
| $\rho$ | 0.014 | $\tilde{b}_1$ | 0.552 | $b_2$ | 1.035 | $b_3$ | 4.14 |

**Table 3:** Regional interaction parameter variation in secondary tauopathy

| Brain region ID and modified $b_3$ value | | | |
| --- | --- | --- | --- |
| Pars Opercularis | 7.452 | Rostral middle frontal gyrus | 6.707 |
| Superior frontal gyrus | 7.452 | Caudal middle frontal gyrus | 7.452 |
| Precentral gyrus | 5.589 | Postcentral gyrus | 3.726 |
| Lateral orbitofrontal cortex | 6.486 | Medial orbitofrontal cortex | 6.486 |
| Pars triangularis | 5.520e-6 | Rostral anterior cingulate | 6.210e-6 |
| Posterior cingulate cortex | 3.45 | Inferior temporal cortex | 13.11 |
| Middle temporal gyrus | 11.04 | Superior temporal sulcus | 8.97 |
| Superior temporal gyrus | 8.28 | Superior parietal lobule | 12.42 |
| Cuneus | 13.8 | Pericalcarine cortex | 13.8 |
| Inferior parietal lobule | 11.73 | Lateral occipital sulcus | 15.18 |
| Lingual gyrus | 13.8 | Fusiform gyrus | 7.59 |
| Parahippocampal gyrus | 11.04 | Temporal pole | 1.104e-5 |

**Table 4:** Primary tauopathy regions and parameters

| Brain region | $b_2$ | $b_3$ | Brain region | $b_2$ | $b_3$ |
| --- | --- | --- | --- | --- | --- |
| Entorhinal cortex | 3.125 | 1.104e-5 | Putamen | 3.795 | 3.795 |
| Pallidum | 2.76 | 2.76 | Precuneus | 3.105 | 3.105 |
| Locus coeruleus | 1.38 | 1.38 |  |  |  |

**Table 5:** Primary tauopathy model parameters

| $A\beta$ system parameters | τP system parameters |
| --- | --- |
| $a_0 = 0.75$ | $b_0 = 0.5$ |
| $a_1 = 1.0$ | $b_1 = 1.0$ |
| $a_2 = 1.0$ | $b_2 = 1.0$ |
| $\tilde{a}_1 = 0.6$ | $\tilde{b}_1 = 0.4$ |
| Coupling parameter: | $b_3 = 1.0$ |

##### Synthetic domain

We set all nodes to the healthy state (*u, ũ, v*, 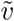) = (0.75, 0, 0.5, 0) and perturb the initial condition of the left-hand nodes 0 *≤ x ≤* 4 by adding a 5% concentration (*ũ* = 0.05) of toxic A*β*. We perturb the initial condition of the right-hand nodes 25 *≤ x ≤* 29 by adding a 5% concentration 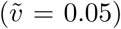 of toxic *τ* P. As expected, we see the toxic A*β* concentration achieve the theoretical maximum, permitted by the parameters, of *ũ* = 0.25 while toxic *τ* P first achieves the maximum associated with 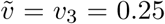 and, upon mixing with A*β*, achieves the fully toxic state value 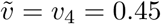. The color scale of Figure 7 was chosen to accentuate the interaction.

**Figure 7:**
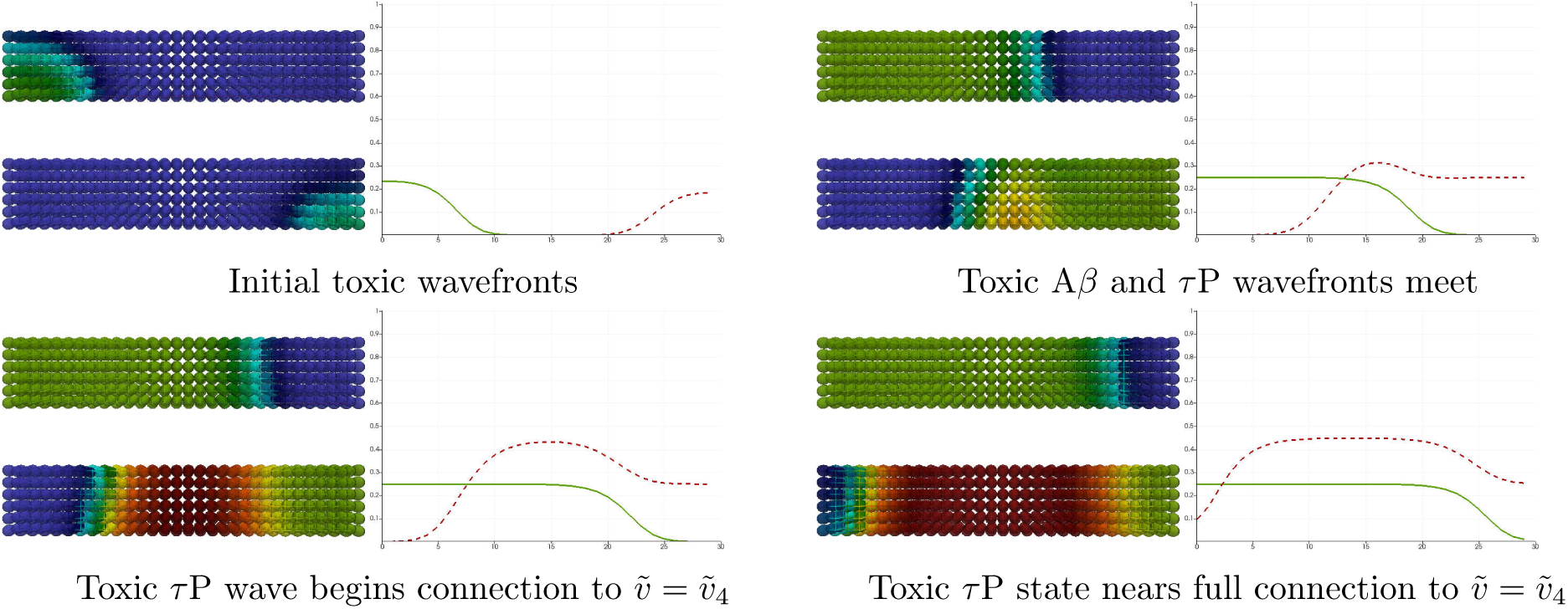
Front propagation in primary tauopathy; synthetic rectangular domain. Each subfigure consists of a toxic A*β* concentration distribution (top left), toxic *τ* P concentration distribution (bottom left) and a plot (solid line: A*β*, dashed line: *τ* P) of the concentration level on the *x* −axis. Dark blue indicates the minimum concentration of *c* = 0.0 while bright red indicates the maximum of *c* = 0.5. See Figure 5 for a comparison to theory. (See also: supplementary movie S1)

##### Brain connectome

Simulation of disease front propagation was then carried out using the physiological connectome of Figure 2. The seeding sites selected for toxic A*β* and toxic *τ* P are the right supramarginal gyrus and left supramarginal gyrus respectively; these seeding sites provide a direct analogy, when the brain connectome is viewed from the frontal lobe, with Figure 7. Figure 8 depicts time instances qualitatively reflecting, in one-to-one correspondence, the stages of the synthetic domain computation of Figure 7. A horizontal slice, at the plane of the supramarginal gyri, of the brain connectome is used to maximally expose the front propagation dynamics. The impact of brain connectome cross-connectivity is evident in the stages depicted in Figure 8. In particular, when the A*β* and *τ* P wavefronts first meet they do so in several locations. This is due to the left-right hemispheric connectivity; both direct nodal connectivity and vis-a-vis propagation in the coronal plane.

**Figure 8:**
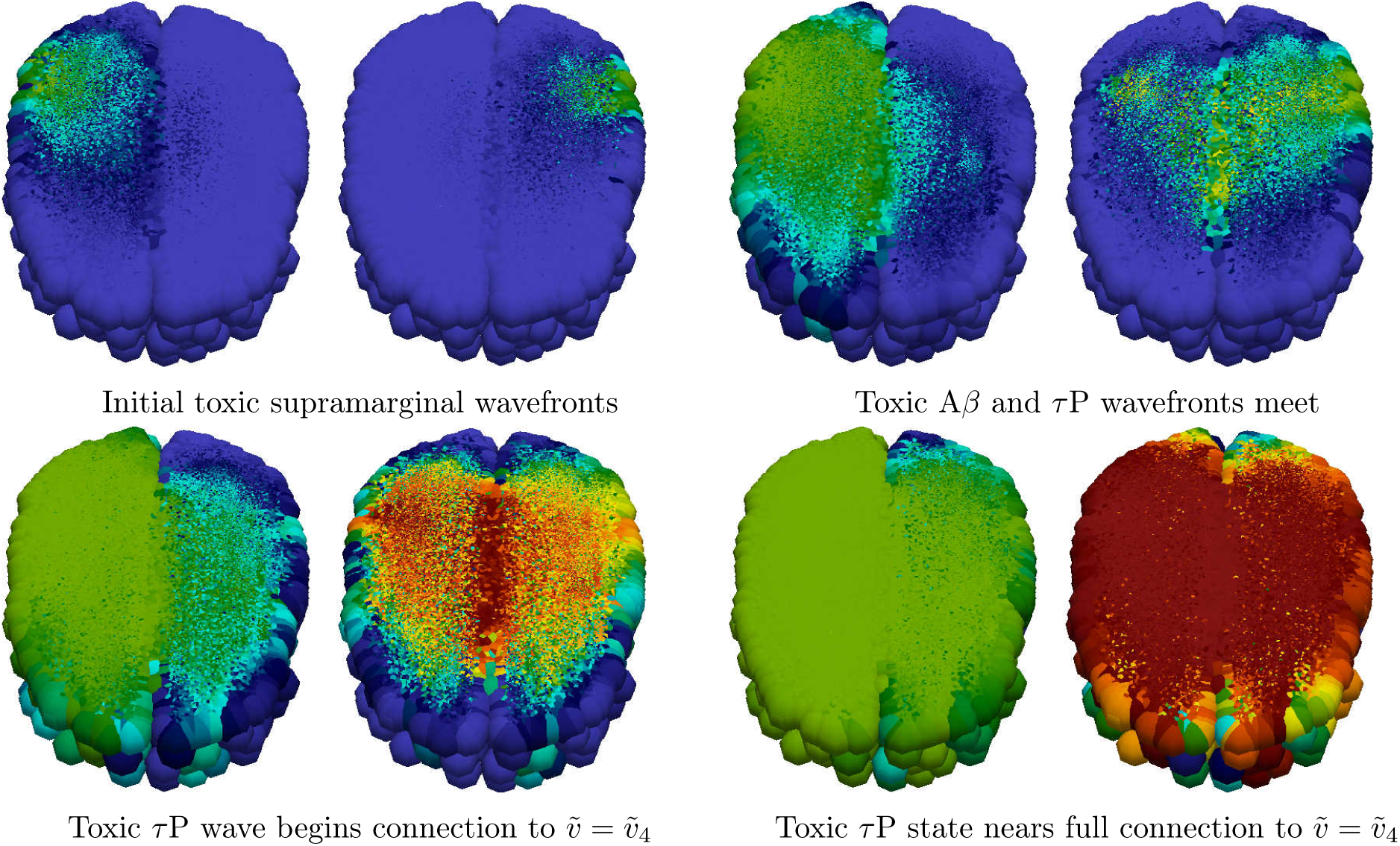
Front propagation in primary tauopathy; brain connectome. Each subfigure consists of a toxic A*β* concentration distribution (subfigure left) besides a toxic *τ* P concentration distribution (subfigure right). Dark blue indicates the minimum concentration of *c* = 0.0 while bright red indicates the maximum of *c* = 0.5. (See also: supplementary movie S2 and supplementary data S9)

#### 4.1.2 Secondary tauopathy

##### Synthetic domain

The parameters for the at-risk secondary tauopathy patient are those of Table 5 with two exceptions; first, as usual for secondary tauopathy, we take *b*_2_ = 0.75 and second we take *b*_3_ = 3.0. We have increased *b*_3_ to facilitate the comparison with Figure 6. As discussed in Section A.2, see also Figure 4, secondary tauopathy consists of all stationary states except for the toxic *τ* P–healthy A*β* state; i.e. (*u*_3_, *ũ*_3_, *v*_3_, 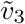) is not included. The stationary point (*u*_4_, *ũ*_4_, *v*_4_, 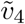) depends on *b*_3_; with the parameters above we have

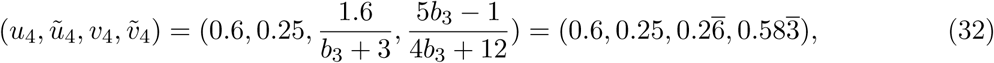

while the other two secondary tauopathy stationary points, c.f. (12)-(13), coincide with their values for primary tauopathy. The initial value at all nodes are first set to the healthy state. A 5% perturbation in concentration is then added to the toxic A*β* initial value for the nodes 0 *≤ x ≤* 4 and a perturbation of 1 *×* 10^−9^%, i.e. 1 *×* 10^−11^, is added to the toxic *τ* P initial value for the nodes 0 *≤ x ≤* 14. As expected: the initial toxic A*β* wavefront achieves its theoretical maximum of *ũ* = 0.25; c.f. Figure 9 vs. Figure 6. The toxic *τ* P wave takes on detectable concentration levels at the point when the A*β* wave reaches the halfway mark in the rectangular domain. The toxic *τ* P state connects, immediately, to the theoretical maximum of the toxic *τ* P–toxic A*β* stationary state value of 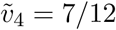 and quickly proceeds to catch up to the A*β* wavefront.

**Figure 9:**
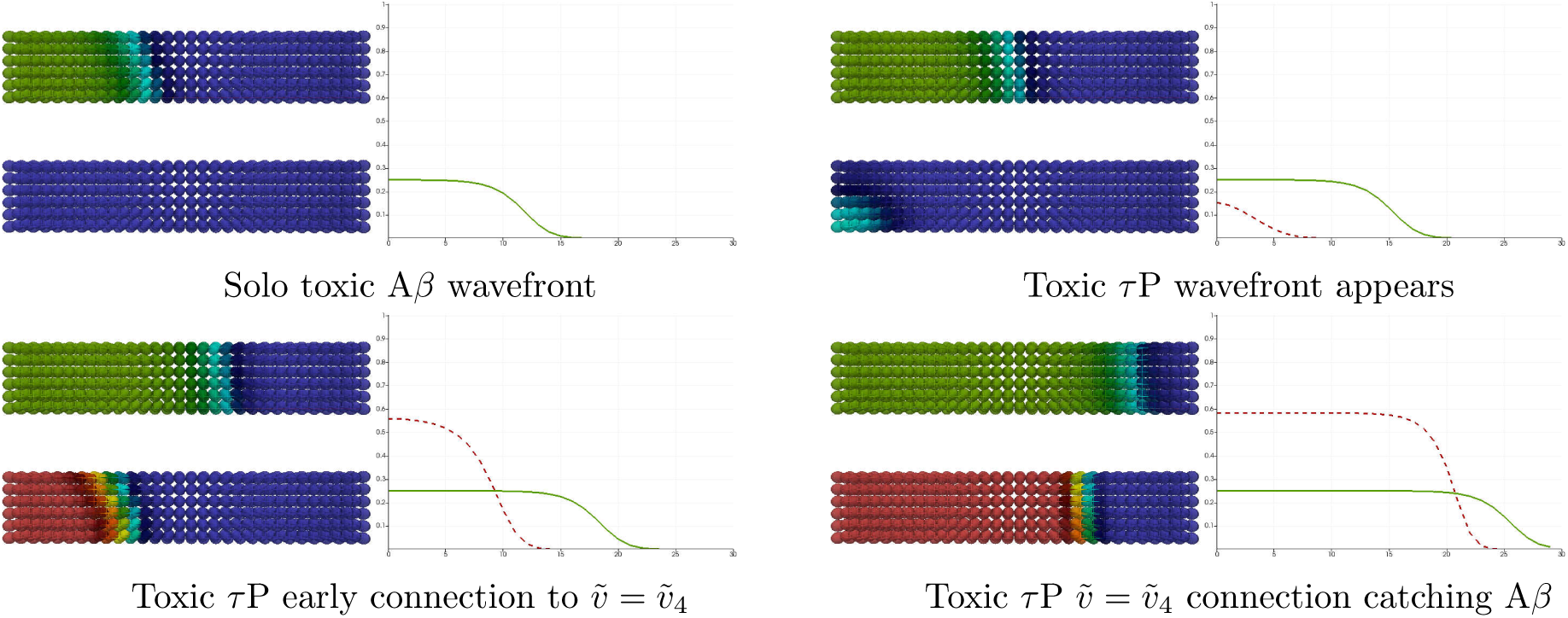
Front propagation in secondary tauopathy; rectangular domain. Each subfigure consists of a toxic A*β* concentration distribution (top left), toxic *τ* P concentration distribution (bottom left) and a plot (solid line: A*β*, dashed line: *τ* P) of the concentration level on the *x* −axis. Dark blue indicates the minimum concentration of *c* = 0.0 while bright red indicates the maximum of *c* = 0.5 for toxic A*β* and 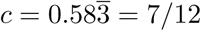 for toxic *τ* P. See Figure 6 for a comparison to theory. (See also: supplementary movie S3)

We tested the time of appearance and saturation of the toxic *τ* P wave front as a function of the interaction parameter *b*_3_. Plots for four values of *b*_3_ are shown in Figure 10 where the y-axis signifies the maximal toxic *τ* P concentration obtained, over all nodes, with respect to the maximum concentration for that value of *b*_3_ (c.f. (32)). Figure 10 highlights the important, and patient-specific, role that *b*_3_ may play in further efforts to deploy (6)-(9) for the modeling of Alzheimer’s disease. In particular values of *b*_3_ *≈* 1 do lead to the development of tauopathy; however, this development emerges significantly later than for higher values of this interaction parameter. Clinically, such a value of *b*_3_ could correspond to a patient who, at the time of death, presents significant amyloid plaques but negligible, or undetectable, levels of neurofibrillary tau tangles.

**Figure 10:**
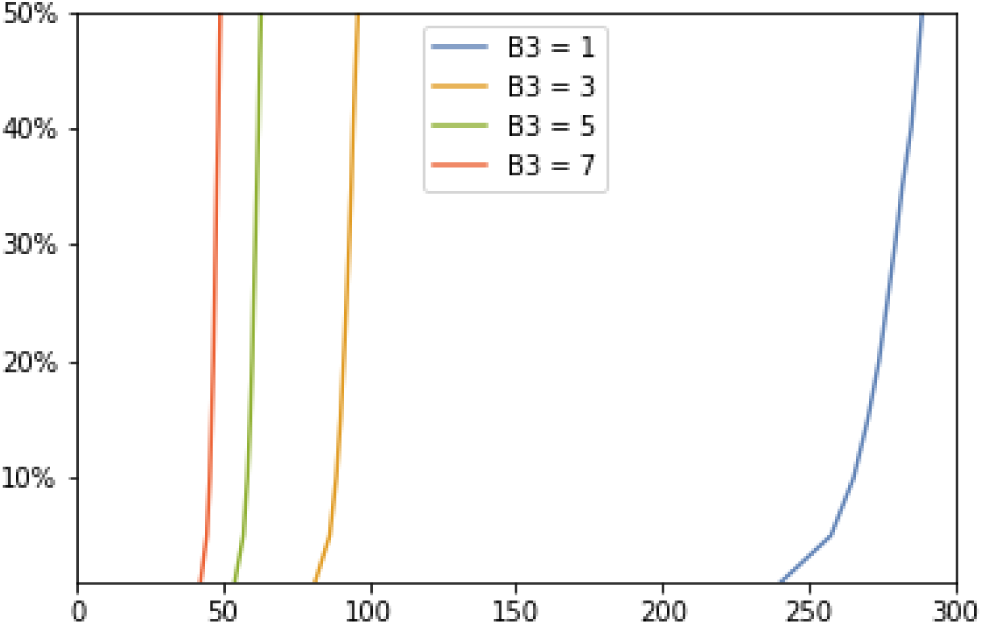
Saturation % (*y*-axis) vs Simulation time (*x*-axis)

##### Brain connectome

We also simulated secondary tauopathy dynamics on the physiological brain connectome of Figure 2. A 5% toxic A*β* perturbation from the healthy state was seeded at the site of the left supramarginal gyrus; all nodes of the left hemisphere were then seeded with an additional 1 *×* 10^−9^% concentration of toxic *τ* P. Snapshots of the evolution is shown in Figure 11. As indicated above we have *b*_3_ = 3 for comparison with Figures 9 and 6. A detail of particular interest is that, even though the entire left hemisphere was seeded uniformly with toxic *τ* P, the toxic *τ* P wave follows the same anisotropic infection pathway, from the left supramarginal gyrus, as the toxic A*β* front propagation. This implies that latent development of tauopathy, in this regime, is heavily influenced by A*β* pathology history.

**Figure 11:**
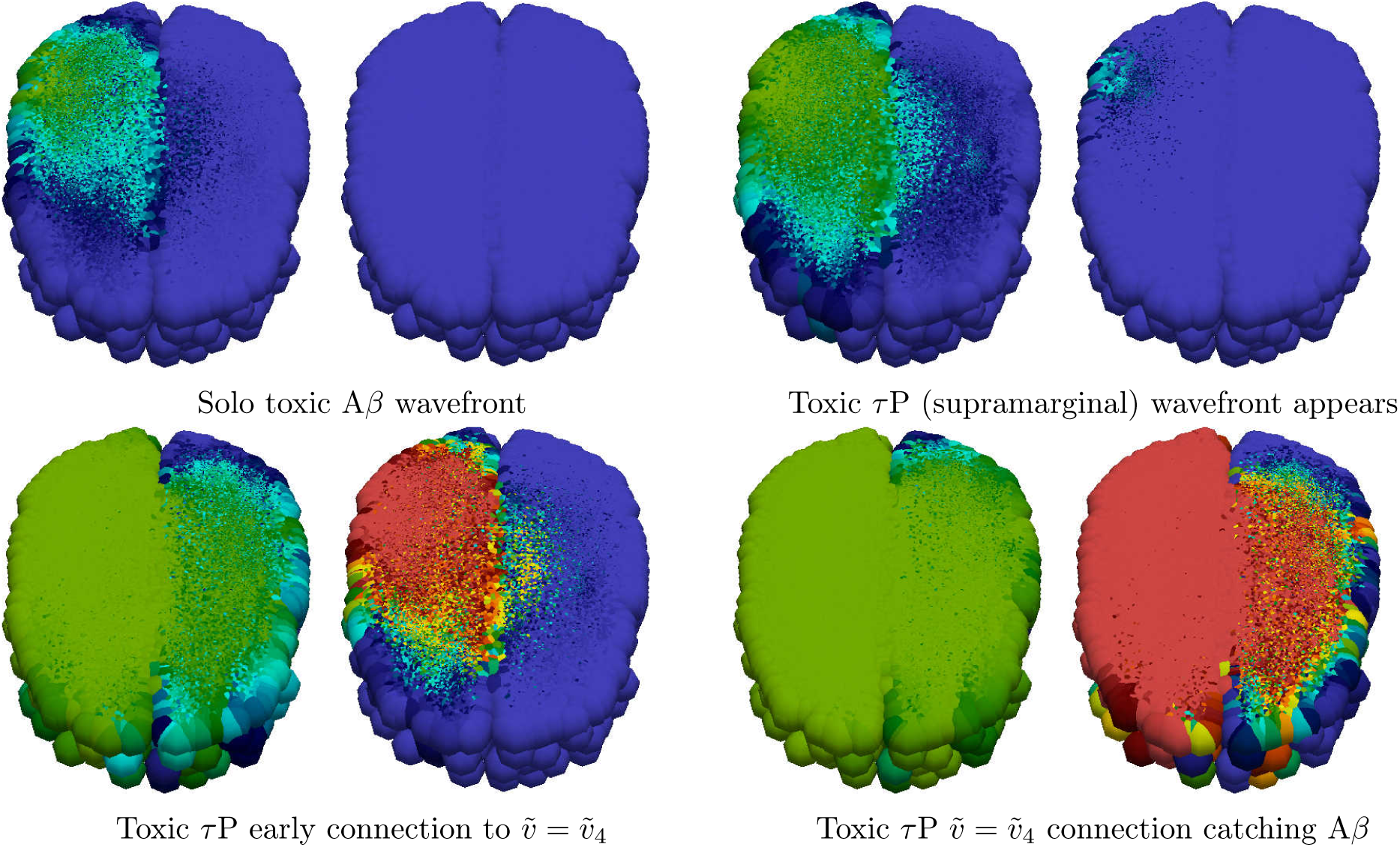
Front propagation in secondary tauopathy; brain connectome. Each subfigure consists of a toxic A*β* concentration distribution (subfigure left) besides a toxic *τ* P concentration distribution (subfigure right). Dark blue indicates the minimum concentration of *c* = 0.0 while bright red indicates the maximum of *c* = 0.5 for toxic A*β* and 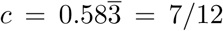 for toxic *τ* P. (See also: supplementary movie S4 and supplementary data S9)

**Figure 12:**
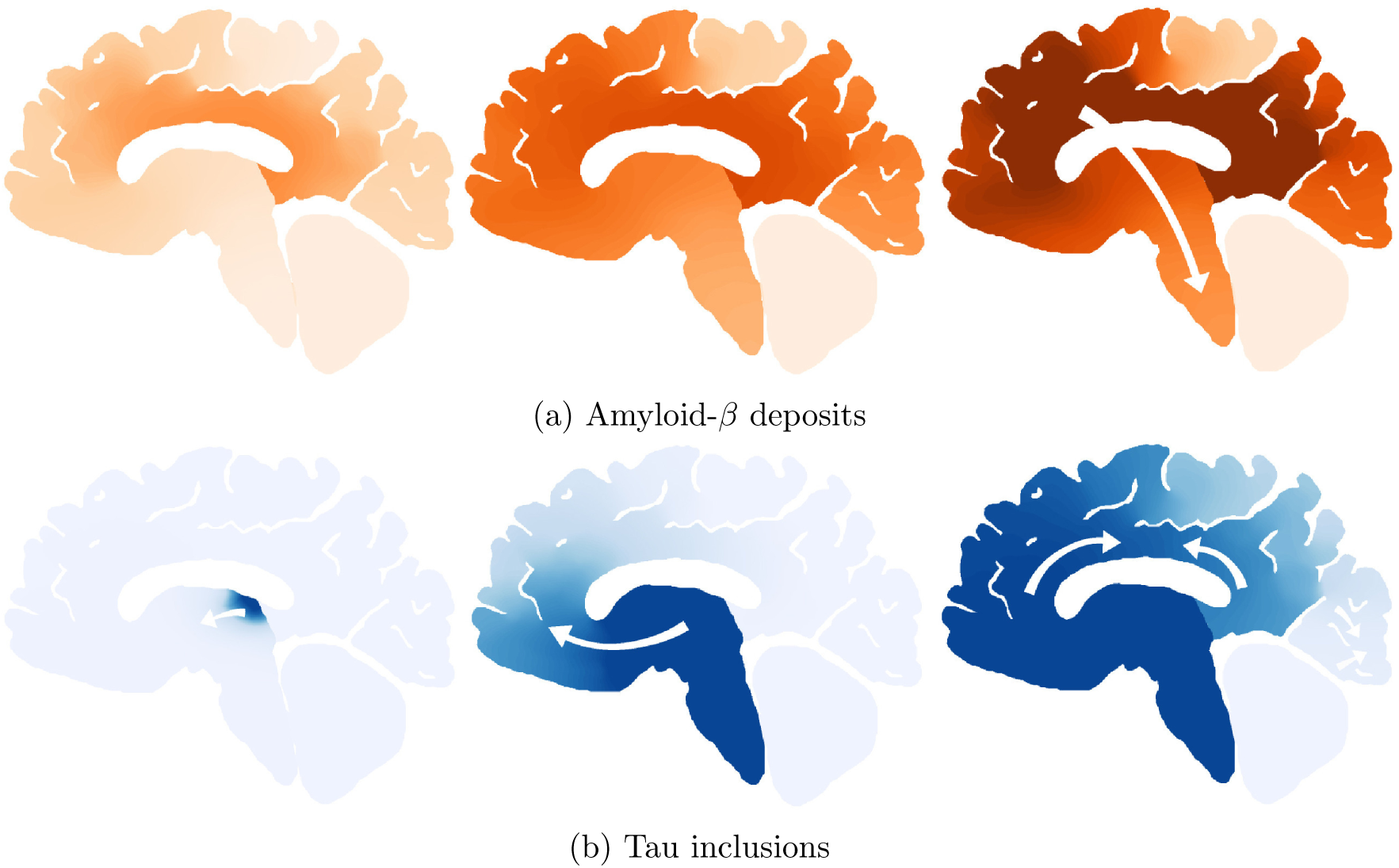
Characteristic progression of of A*β* and *τ* P lesions

## 5 Application to neurodegenerative disease modeling

We have shown in the previous section that the overall phenomenology obtained from the dynamic evolution of the continuous model in one-dimension is recovered within the discrete network setting. We can therefore use the network model and our primary classification to study the interaction of proteins in the brain.

Here, we apply (6)-(9) to a computational case inspired by Alzheimer’s disease. In particular we consider seeding sites, for toxic A*β* and toxic *τ* P, commensurate with [11, 66, 67, 47] Alzheimer’s disease staging. Alzheimer’s disease is a complex multiscale phenomena; a uniform parameter regime, throughout all brain regions, is unlikely to accurately reflect a patient’s real disease progression. Nevertheless, for this early investigation, we will consider the simple uniform parameters, of the model’s primary and secondary tauopathy regimes, as discussed in Section 4. In addition we briefly consider the evolution of the coupled neuronal damage term, given by (10), and the effect of the coefficients therein. We shall also select the diffusion constants, *ρ* of (5), to be unity for (6)-(9).

### 5.1 A simplified model of Alzheimer’s disease proteopathy

Alzheimer’s associated amyloid deposition begins [18, 47, 66, 67] in the temporobasal and frontomedial regions. Tau staging, in Alzheimer’s disease, follows the Braak tau pathway [11] and begins in the locus coeruleus and transentorhinal layer [18, 47, 67]. These seeding sites, used throughout this section, are shown in Figure 13. The temporobasal and frontomedial regions for toxic A*β* seeding are highlighted in red on the left while the locus coeruleus (in the brain stem) and transentorhinal associated regions, for toxic *τ* P staging, are highlighted red on the right. In sections 5.1.1 and 5.1.2 we will investigate primary and secondary tauopathy, respectively, on the whole brain connectome with globally-constant synthetic parameters. We will observe several characteristic traits of these modalities and also note the similarity between these pure states and to a qualitative three-stage progression [18] of protein lesions, typical of Alzheimer’s disease, as inferred from post-mortem analyses. In section 5.1.3 we consider the case of mixed regional modalities; i.e. a mixture of primary and secondary tauopathy connectome regions. We illustrate that the model can manifest canonical features of positron emission tomography (PET) SUVR intensities characteristic of Alzheimer’s disease (c.f. for instance [68, 69]). In particular: we will compare the results of a mixed-mode simulation with a cross sectional Alzheimer’s patient cohort dataset procured from the Alzheimer’s Disease Neuroimaging Initiative (ADNI) database.

**Figure 13:**
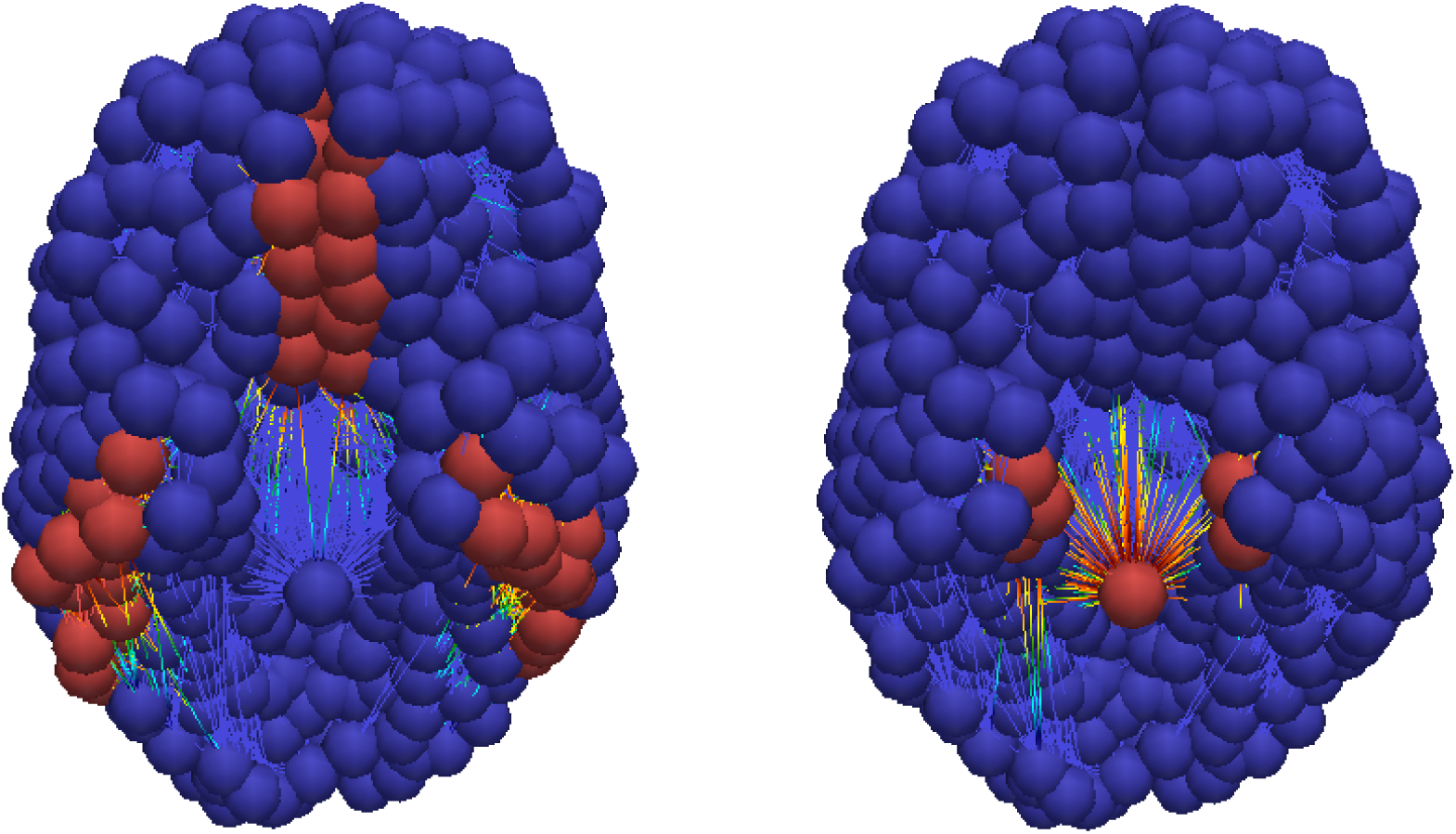
Toxic A*β* (left) and toxic *τ* P (right) seeding sites

#### 5.1.1 Primary tauopathy

All nodes in the connectome were first set to the healthy, but susceptible, primary tauopathy patient state; c.f. (34). The temporobasal and frontomedial A*β* seeding sites, consisting of fifty-three nodes, were each seeded with a toxic amyloid concentration of 0.189%; thus the brain-wide toxic A*β* concentration represents a 1% concentration deviation from healthy. Similarly, the locus coeruleus and transentorhinal nodes were seeded with an aggregate perturbation of 1% toxic *τ* P.

Figure (14a) shows the average brain-wide concentration for all four protein populations for the primary tauopathy patient (c.f. Table 5) with interaction term *b*_3_ = 1. As we observed previously, in (32), the value of *b*_3_ directly informs the saturation *τ* P concentration, of 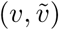, for the disease. Figure 14b shows the evolution of the toxic *τ* P burden for various *b*_3_.

**Figure 14:**
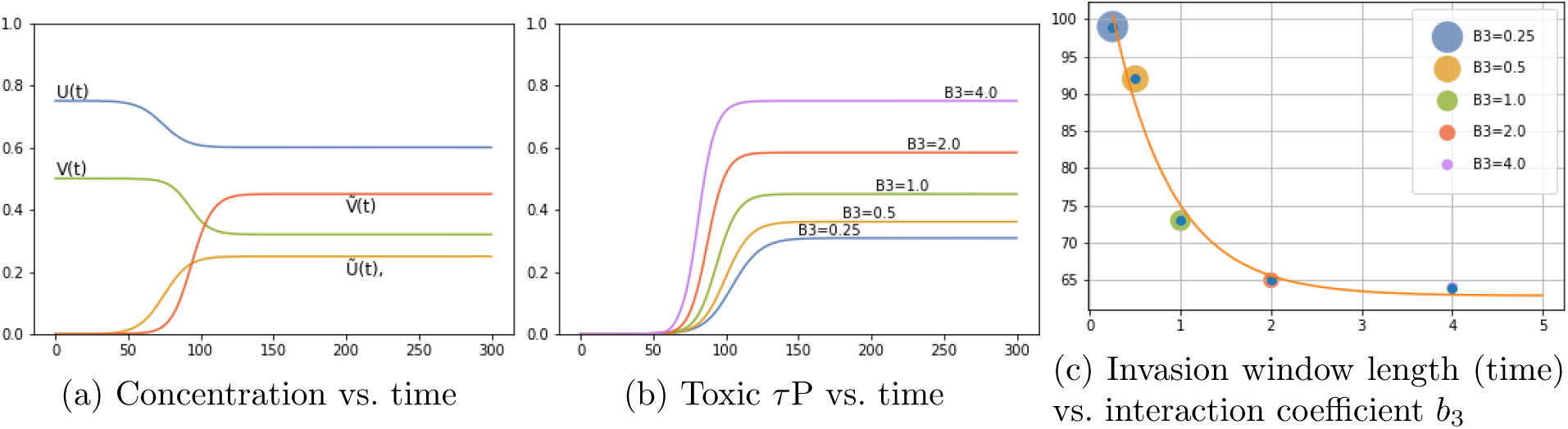
Protein-protein interaction in primary tauopathy

For each value of *b*_3_ the toxic *τ* P invasion window was computed as the difference in time between the appearance of a global 1% toxic *τ* P concentration to the simulation time where the maximum 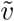 was reached. We performed a least squares fit and found that the invasion window, for primary tauopathy, decreases exponentially with an increase in coupling strength (*b*_3_) between toxic A*β* and toxic *τ* P. Figure 14c shows the result. This result suggests that the dynamics of toxic protein evolution is highly sensitive to the coupling between A*β* and *τ* P: Toxic A*β* accelerates, in a nonlinear fashion, the way toxic *τ* P emerges across the brain. Acceleration of toxic *τ* P progression due to the presence of toxic A*β* has also been observed in mouse models of Alzheimer’s disease [27]. Consulting longitudinal tau PET studies, in combination with amyloid-beta data from a public database, could provide an estimation of *b*_3_ in the primary tauopathy model.

The toxic load progression of the susceptible primary tauopathy patient is shown in Figure 15 at five equidistant time points throughout the invasion window. To facilitate a comparison with Figure 12: a sagittal view of the progression, of each toxic agent, is presented; directly below is an opacity-exaggerated view wherein regional opacity is proportional to the agent’s regional toxic load. Comparison with Figure 12 suggests reasonable qualitative agreement; thus warranting further study of physically relevant parameters with a view towards real clinical applications.

#### 5.1.2 Secondary tauopathy

All nodes were set to the healthy, but susceptible, patient state corresponding to the susceptible secondary tauopathy patient parameters (Table 5 with *b*_2_ = 0.75). In addition, for a baseline secondary tauopathy case, we follow Section 4.1 and select the interaction parameter of *b*_3_ = 3.0; the fully invaded secondary tauopathy state values are therefore (32). Seeding patterns for both A*β* and *τ* P are identical to the case of primary tauopathy discussed above.

Figure (16a) shows the average brain-wide concentration for all four protein populations of the secondary tauopathy patient with baseline interaction term *b*_3_ = 3. As in the case of primary tauopathy we investigate the effect of *b*_3_ on toxic load and invasion window by considering a value range four times smaller to four times larger than the baseline *b*_3_ = 3 case. Toxic load curves are shown in Figure 16b while invasion windows are shown in Figure 16c.

Interestingly, we see distinct differences in comparison with the primary tauopathy case (c.f. Figures 14a–14c). More specifically, in primary tauopathy it is evident (Figure 14b) that the disease onset is only slightly affected by varying the interaction parameter *b*_3_; for secondary tauopathy, in contrast, *b*_3_ has a profound effect on disease onset latency. Moreover, the invasion window variation with *b*_3_ for secondary tauopathy is more complex than that of primary tauopathy. Figure 14c shows that the invasion window duration initially decreases exponentially with *b*_3_ but then appears to increase logarithmically for *b*_3_ *≥* 3.

Analyzing the invasion window start time and end time separately shows a clear, but separate, exponential decay pattern versus *b*_3_. Figure 17 shows the least-squares exponential fit to the invasion start and end times. As in the primary tauopathy case we now consider characteristic toxic load progression for secondary tauopathy. The A*β* progression is identical to that shown in Figure 15 (top two rows). This is expected as only the *τ* P portion of the system has been modified with respect to the primary tauopathy regime; see Section A.2. The *τ* P secondary tauopathy progression is shown, in Figure 18, at equally spaced simulation times through the invasion window. Qualitatively, the progression of secondary tauopathy also reflects the characteristic post-mortem progression of Figure 12.

**Figure 15:**
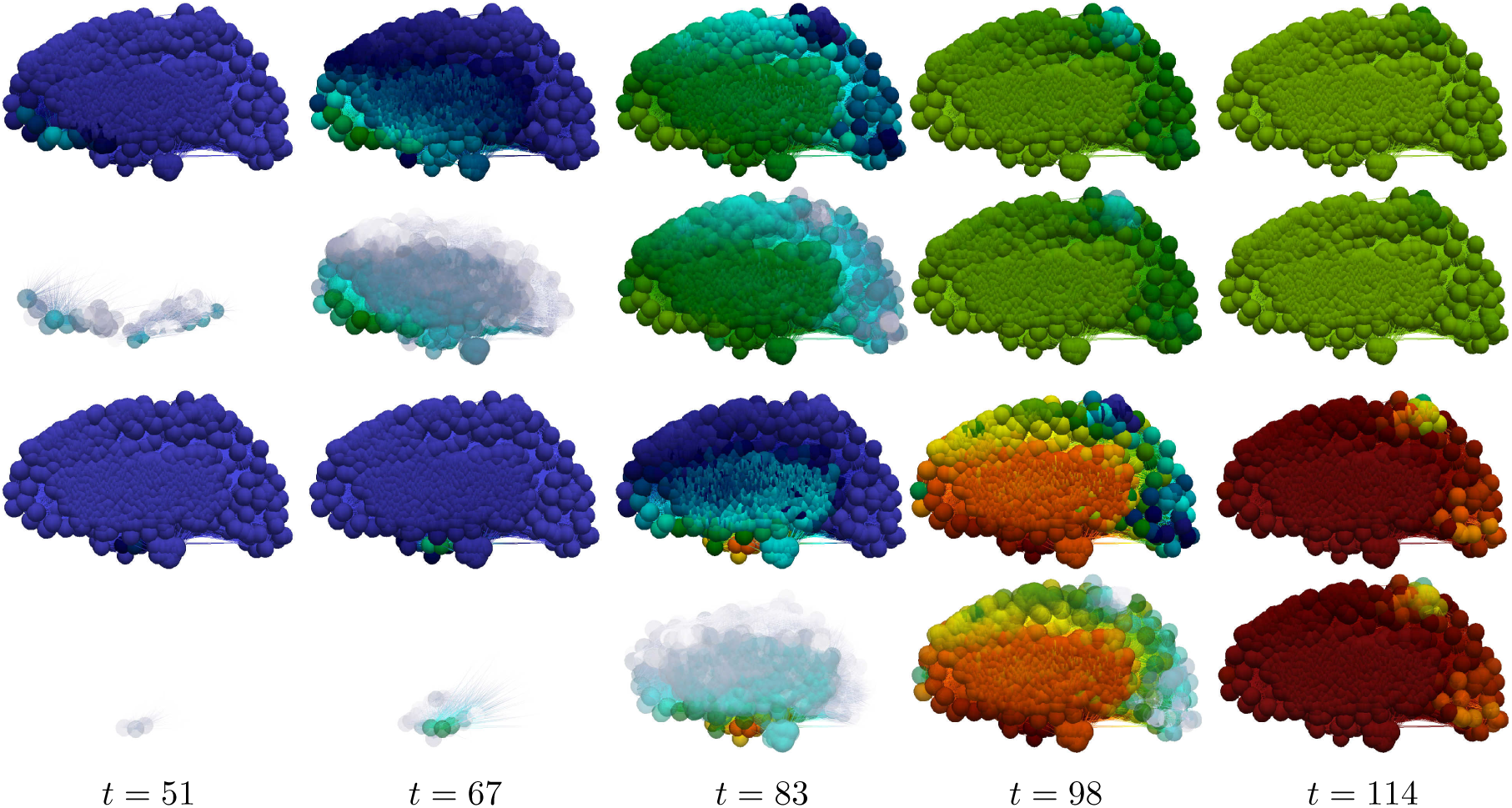
Toxic proteopathy progression dynamics in the primary tauopathy patient. Toxic A*β* (top row) and opacity exaggerated toxic A*β* progression (second row); Toxic *τ* P (third row) and opacity exaggerated toxic *τ* P progression (last row). Color scale is identical to Figure 7. (See also: supplementary movie S5, supplementary movie S6 and supplementary data S9)

**Figure 16:**
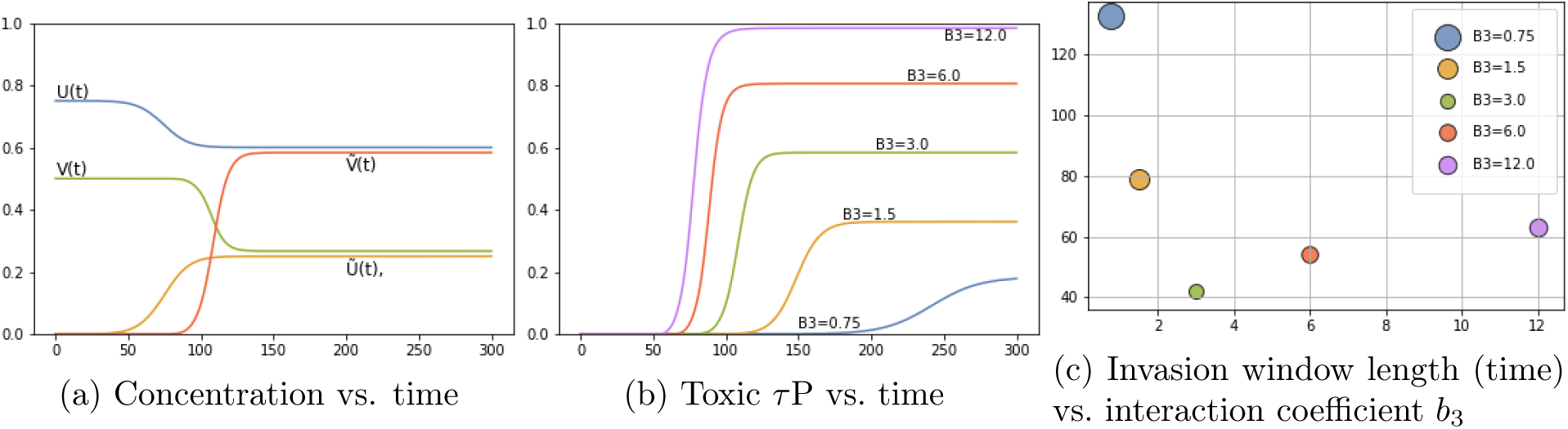
Protein-protein interaction in secondary tauopathy

**Figure 17:**
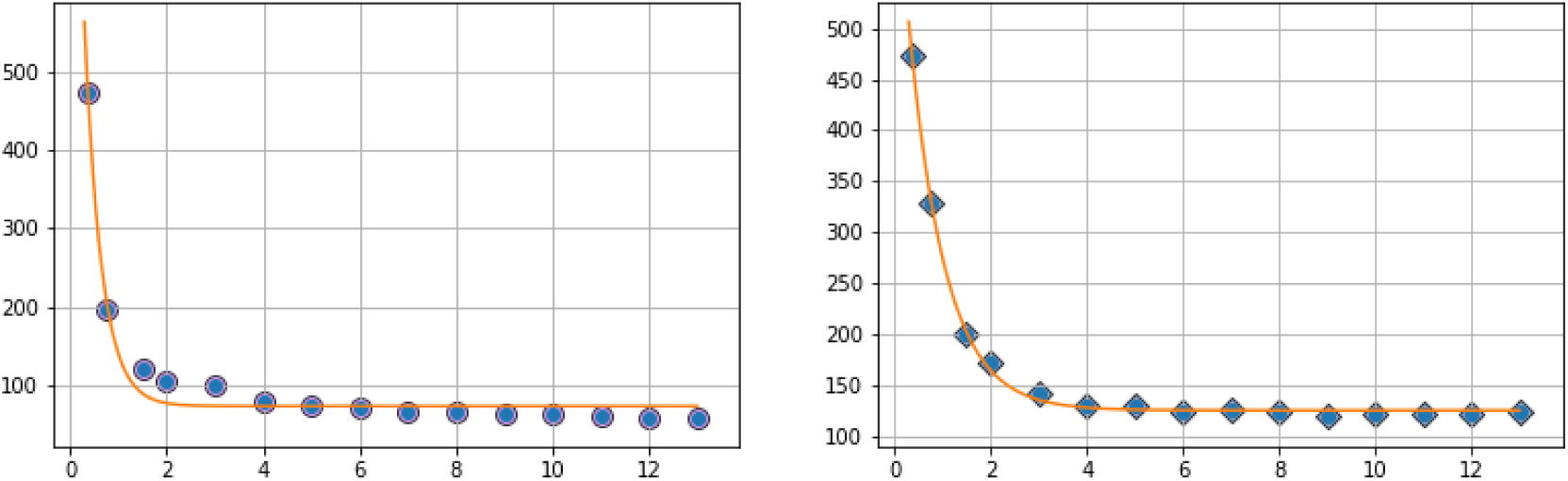
Invasion starting (left) and ending (right) time vs. *b*_3_

**Figure 18:**
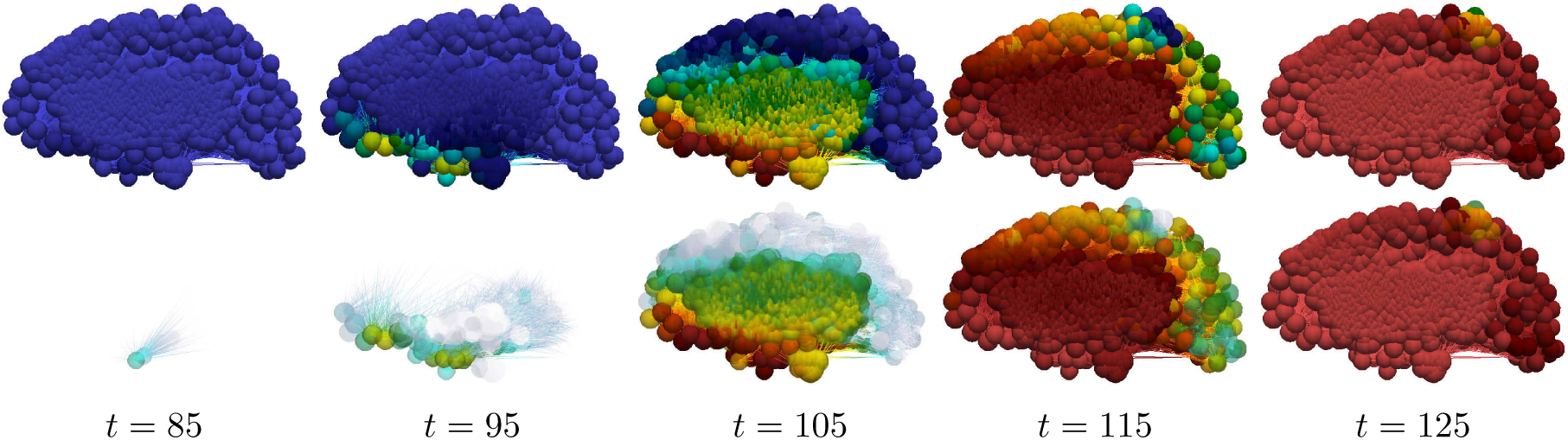
Toxic *τ* P progression dynamics in the secondary tauopathy patient. Toxic *τ* P (first row) and opacity exaggerated toxic *τ* P progression (second row). Color scale is identical to the *τ* P case of Figure 9. (See also: supplementary movie S5, supplementary movie S7 and supplementary data S9)

#### 5.1.3 A mixed model comparison to Alzheimer’s diseased patient data

Sections 5.1.1 and 5.1.2 have focused, respectively, on the general features of the modalities of primary versus secondary tauopathy; illustrated with synthetic, globally constant parameters. We have observed several interesting facets of these two disease states. For instance, regions in a state of primary tauopathy (Sec. 5.1.1) can develop A*β* and *τ* P proteopathy separately; the A*β* interaction parameter, *b*_3_, does not alter the onset of tauopathy but does modulate the regional concentration. Conversely, in secondary tauopathy (Sec. 5.1.2), the presence of A*β* pathology is necessary for *τ* P pathology and the interaction parameter, *b*_3_, modulates both the latency and the intensity of the regional pathology. We have also seen that proteopathy progression in pure models, e.g. where all regions have the same primary or secondary tauopathy parameters, bears a notable resemblance (Fig. 12) to post-mortem progression of protein lesions [18]. However, PET imaging studies of A*β* and *τ* P radiotracer uptake tell a more nuanced story. For instance, in Alzheimer’s disease the distribution of ([^18^F]flortaucipir and [^18^F]THK-5117, among others) PET-*τ* P SUVR intensities are distinctly biased [68, 69] towards the temporal and parietal regions of the brain; a feature that we do not see in Figure 15 or Figure 18.

In order to demonstrate that the model of (6)-(9) can reproduce salient features of A*β* and *τ* P SUVR uptake in patients diagnosed with Alzheimer’s disease: we now compare a mixed-modality simulation with a cross-sectional study of Alzheimer’s disease patient data. Sample data for model comparison was procured from the the Alzheimer’s Disease Neuroimaging Initiative (ADNI) database. We first queried the ADNI database to locate AD-diagnosed subjects between the age of 70 and 90 who had at least one *τ* P PET (18F-AV1451, flortaucipir) scan. The returned results consisted of 41 patients. These initial patient candidate IDs were then checked for a structural T1 weighted MRI within a maximum of one year of a tau PET scan; patients without any sMRI, those with a poor quality sMRI, or those without an sMRI within a year of the tau PET scan were discarded. The resulting cohort consisted of 38 patients (25 male and 13 female) with mean age 78.4, a standard deviation of 5.14 years, and a male-to-female ratio of 1.92. Patients who met the age, *τ* P PET scan, and acceptable quality sMRI within one year of the PET scan were not abundant in the ADNI database. Due to this the *τ* P group was selected, first, to maximise the number of candidates in the group.

The ADNI database contains quite a generous number of patients with A*β* (18F-AV45, florbetapir) PET scans. We next queried the database to locate AD-diagnosed subjects between 70 and 90 who had an A*β* (18F-AV45) PET scan in addition to a structural T1 weighted MRI within one year. The result of this search was in excess of 100 unique patient IDs; from these results we selected an initial candidate group of 82 unique patients IDs. The 82-candidate group was further pruned to create a list of 48 subjects whose age and sex characteristics closely resembled that of the *τ* P PET group. Finally, the 48 candidate A*β* PET group was narrowed down: first, subjects with an unacceptable or low-resolution sMRI were removed. We then removed the minimum number of candidates required to provide as close a match as possible to the mean age, standard deviation and male-to-female ratio of the *τ* P PET group. The resulting AD cohort for AB consisted of 42 patients with mean age 78.4, a standard deviation of 5.1 years, and a male-to-female ratio of 2.0. The two groups are succinctly summarised in Table 1.

Patient data was then processed through a semi-automated, scripted software pipeline for general connectome-graph based imaging and analysis of clinical patient data. Each of the 160 patient images, the PET and sMRI scan for each patient, were first manually analysed using version 12 of the Statistical Parameteric Mapping [70] (SPM) software; the origin of the image was set to coincide with the anterior commissure. Next, the sMRI images for each patient were pre-processed for connectome-graph visualization. The SPM software was used, on each patient sMRI, to perform a unified segmentation procedure [70]. The unified segmentation procedure identifies grey matter, white matter, cerebrospinal fluid, skull, and exterior regions. Following this, the grey and white matter segmentations served as input for spatial normalisation using the DARTEL [71] toolbox; the outputs of which were a composite template for the A*β* patient group and a separate composite template for the *τ* P patient group. All patient grey matter images were then normalised to MNI-152 space using their group-specific optimised DARTEL template.

The next step of the pipeline is to treat the PET images for both the A*β* and *τ* P groups. This step relies on the fact that we have already manually relocated the origin of the PET images to the approximate visual location of the anterior commissure as mentioned above. The first new step for this portion of the pipeline is to use the SPM software to co-register the PET images to their sMRI counterparts. This co-registration step is the genesis of the original data procurement requirement that an sMRI scan is conducted no later than one year beyond the PET image acquisition date. The coregistered PET images are then spatially normalised using the DARTEL template and corresponding subject deformation fields derived from the sMRI pipeline (c.f. above). Finally, SUVR values were computed, using SPM, by means of a whole-cerebellar reference region; the skull was then stripped. A voxel-wise mean, across all subjects, was taken to produce a representative SUVR map of both AD cohorts. This completes the first portion of the connectome-graph based imaging analysis pipeline; the result of this step, for both the A*β* and *τ* P group, is shown in the top row of Fig. 19.

**Figure 19:**
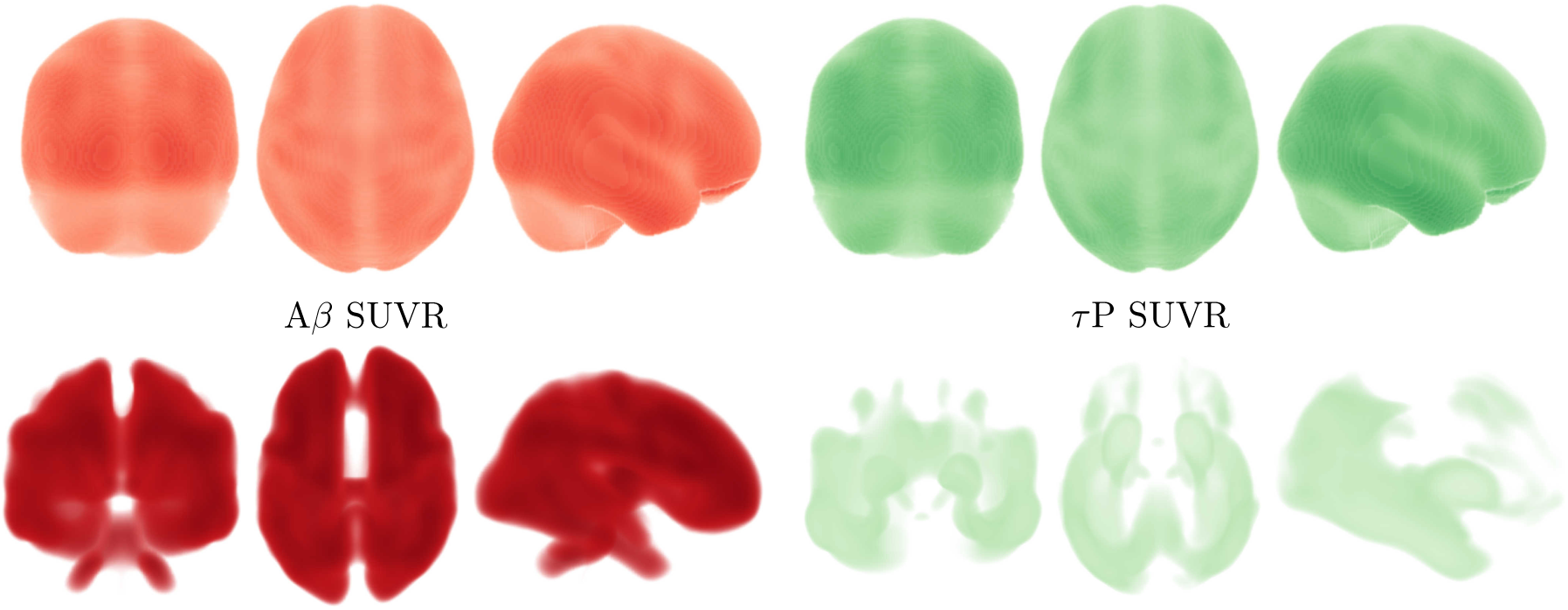
Skull-stripped, cross-sectional Alzheimer’s patient cohort SUVR intensity. Top row: averaged SUVR data is shown. Bottom row: top 30% of SUVR intensities are visible. For both rows: (left side) 18F-AV45 florbetapir A*β* radiotracer SUVR and (right side) 18F-AV-1451 flortaucipir *τ* P radiotracer SUVR. Darker colors correspond to higher SUVR values.

The averaged SUVR data of Fig. 19 (top row) reports a general view of uptake across the whole brain. In order to visualize significant features of the data: the skull-stripped SUVR image volumes, in NIfTI file format, are visualized using Paraview vis-a-vis the NIfTI Paraview plugin. The volume opacity then set so that the top 30% of the SUVR intensity range in the data is visible; c.f. Fig. 19, middle row. Doing so: we immediately see notable features of significance reported in previous radiotracer studies; in particular the familiar [68, 69] temporal and parietal dominance of the *τ* P radiotracer uptake distribution (Fig. 19, middle right) are visible. In order to compare simulation results to the patient data of Fig. 19 we now employ a connectome-graph data visualization software process. This portion of the general pipeline uses functionality from both SPM and the Nilearn [72] Python library. Regional masks were produced using the Lausanne multiresolution atlas [73] parcellation to the MNI ICBM 152 non-linear 6th generation symmetric volume [74]; generating over 1000 distinct masks. The mask volumes were then applied to isolate the SUVR values for each mask in the parcellation and a regional average SUVR was computed. The computed values were then normalized to lie in the interval [0, 1] by dividing all regional SUVR averages by the global maximum average SUVR intensity; the normalized values, along with the MNI-space coordinates of the region’s centroid were recorded as output. The regional normalised SUVR values and the MNI coordinates were then used as input to the Python application programming interface of the Nilearn [72] software. Using the connectome visualization capabilities of Nilearn we rendered this information using a glass brain view with the highest 30% of values shown; see Fig. 20. A comparison with Fig. 19 (bottom row) shows that characteristic PET features associated with Alzheimer’s disease [68, 69] are once more prominent in the connectome view of the top 30% of SUVR intensities.

**Figure 20:**
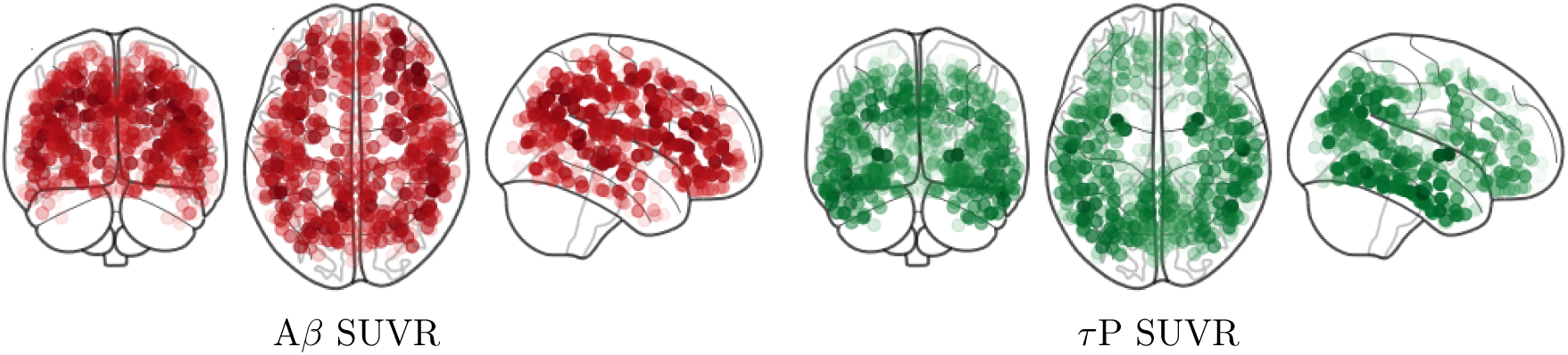
A connectome-graph view of the normalized SUVR for the (left side) 18F-AV45 florbetapir A*β* radiotracer SUVR and (right side) 18F-AV-1451 flortaucipir *τ* P radiotracer SUVR. Highest 30% of connectome regional values are visible. Darker colors correspond to higher SUVR values.

Demonstrating that the mathematical model of (6)-(9) is capable of achieving distributions of toxic A*β* and *τ* P that resemble the PET data of AD patients is a multi-step process. We note that the demonstration endeavored here is illustrative and does not constitute a full validation of the model; it will, however, fully justify that the fitting of real-world data is within the capacity of the model. First, we set all regions in the connectome to a state of secondary tauopathy with the general synthetic parameters given by those in Table 2. The A*β*–*τ* P interaction parameter, *b*_3_, was modified in several regions. All of the modifications to *b*_3_ were symmetric; that is, they were made in both the left and right hemispheres of the corresponding region. The modified interaction parameters for connectome vertices in select secondary tauopathy regions are shown in Table 3. Finally, the connectome vertices in a total of five brain regions, in both hemispheres, were put into a state of primary tauopathy by changing the values of *b*_2_ and *b*_3_ to correspond to states in this regime. The primary tauopathy regions, and their parameters, are listed in Table 4.

The connectome vertex parameters given by Table 2 and regional vertex parameter modifications pursuant to Table 3 and Table 4 describe a mixed-modality mathematical model; the connectome graph contains vertices in a state of primary tauopathy and vertices in a state of secondary tauopathy. The model equations (6)-(9) were solved with the regional parameters, and modifications, described above. Seeding patterns for both A*β* and *τ* P are identical to those of Sections 5.1.1 and 5.1.2. The results of the simulation are shown in Figure 21 at time *t* = 78 in accordance with the mean age of the A*β* and *τ* P cross-sectional study parameters (c.f. Table 1); the highest 30% of values are visible. A comparison of Fig. 19 (bottom row) and Fig. 20 to that of Fig. 21 shows that the model can indeed capture salient characteristics of Alzheimer’s disease proteopathy as indicated by SUVR intensity. Thus, this preliminary result clearly demonstrates that the model can recover primary features of Alzheimer’s disease proteopathy and that more mathematically comprehensive analyses are warranted; for instance, investigations using (variational) Bayesian methods [75, 76] may be compelling for further study of the model alongside patient data for cognitively normal, mildly cognitively impaired, early and late onset Alzheimer’s disease cohorts.

**Figure 21:**
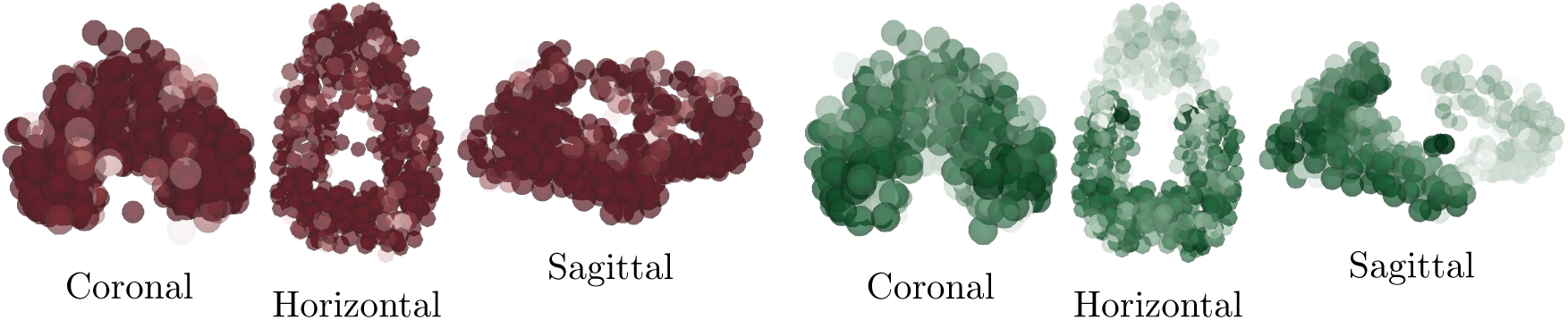
Results of a mixed-modality simulation. (left) Toxic A*β* population and (right) toxic *τ* P population are shown at time *t* = 78. The top 30% of nodal values are visible; darker colors correspond to higher values.

### 5.2 A simple model of local and non-local neuronal damage

This section briefly examines the use a simple measure of neuronal damage with the minimal level of complexity necessary to take into account both local and non-local effects. The intent is to explore the qualitative differences between the primary and secondary tauopathy regimes and the effect of varying: the toxification rate of A*β* on *τ* P; and the rate of aggregation due to nonlocal influence. In Section 2.1, the continuous equations (1) were augmented with a coarse-grained damage model (2). We recall that *q*(**x**, *t*) represents a first-order assessment for neuronal cell body damage vis-a-vis a, potentially variegated, set of coupled mechanisms. These mechanisms are not individually differentiated; however, they are assumed to be correlated with the presence of toxic A*β*, toxic *τ* P or with those mechanisms requiring both (c.f. the discussion surrounding equation (2)).

The damage model (2) has several coefficients: *k*_1_ and *k*_2_ mediate the damaging effect of toxic A*β* and *τ* P respectively. The rate coefficient *k*_3_ reflects damage, such as the rate of neuronal death following over-excitation, resulting from the combined presence of toxic A*β* and toxic *τ* P. Finally, *k*_4_ determines the rate of transneuronal damage propagation; thus reflecting aggregate neuronal death as a result of communication disruption to and from regional neighbors.

In this illustrative example we consider the parameters

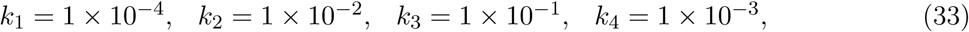

as a baseline from which to begin investigation. These parameters have been chosen to reflect a few clinical observations. First, *k*_1_ is chosen as significantly less than *k*_2_ to reflect the correlation [9, 10, 12, 13] of toxic *τ* P neurofibrillary tangles with various forms of neuronal damage (e.g. intracellular NFT-induced neuron death, atrophy etc). Second, toxic effects of *τ* P are increased in the presence of toxic A*β* [12, 26, 28, 29, 30, 31, 32, 33] thus, *k*_3_ is taken larger than *k*_2_.

As a first point of enquiry: we consider our baseline tauopathy patient parameters, laid out in Section A.2, and vary the deafferentation parameter *k*_4_ across three orders of magnitude from the initial value given in (33). Figures 22a–22b show the results. Note that, in each subfigure, the dashed lines correspond, from left to right, to monotonically decreasing values of *k*_4_; the far left dashed curve is *k*_4_ = 1.0, the next curve to the right is *k*_4_ = 1 *×* 10^−1^, the next is *k*_4_ = 1 *×* 10^−2^, and so forth, down to the final (rightmost) curve corresponding to *k*_4_ = 1 *×* 10^−6^. In both figures the baseline deafferentation curve, *k*_4_ = 1 *×* 10^−3^, is instead solid (and red) for emphasis. Figures 22c–22d show the effect of increasing *b*_3_; we have incremented *b*_3_ by two, from baseline, for each case. As expected an overall increase in toxic *τ* P, 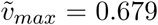 for primary tauopathy and 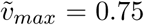 for secondary, is observed with the increase in *b*_3_. However, the limiting behavior of the deafferentation baseline coefficient choice, *k*_4_ = 1 *×* 10^−3^, remains; which justifies our choice of *k*_4_ in (33).

**Figure 22:**
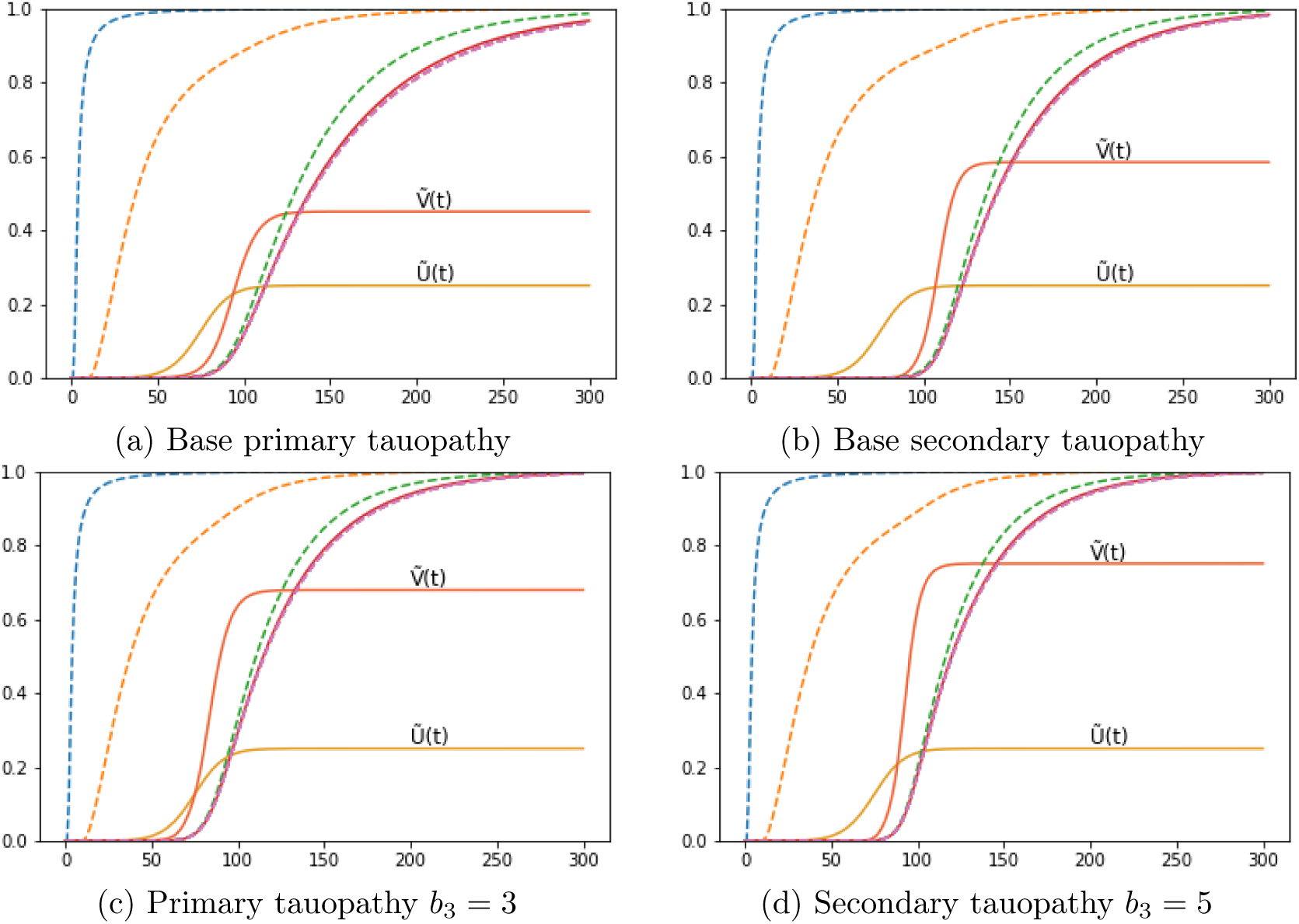
Aggregate damage (dashed; except *k*_4_ = 1 × 10^−3^ solid, red) curves in the base primary (a) and secondary (b) tauopathy patients. Damage with increase toxic protein interaction, b3, in primary (c) and secondary (d) tauopathy.

The staging of the damage is presented in two figures: primary tauopathy in Figure 23 and secondary tauopathy in Figure 24. Each set of figures includes an overhead horizontal plane view in addition to a sagittal view of the right hemisphere. A visualization starting time was selected to coincide with the first visibility of 5% damage, in any nodes, while an ending time was selected such that the damage progression appeared qualitatively equal. Progression times are uniformly spaced within this interval to allow for a direct comparison between the damage distribution within the two regimes. An immediate observation is that a 5% damage detection is latent within the secondary model, starting at *t* = 95, compared to the primary tauopathy paradigm at *t* = 80.

**Figure 23:**
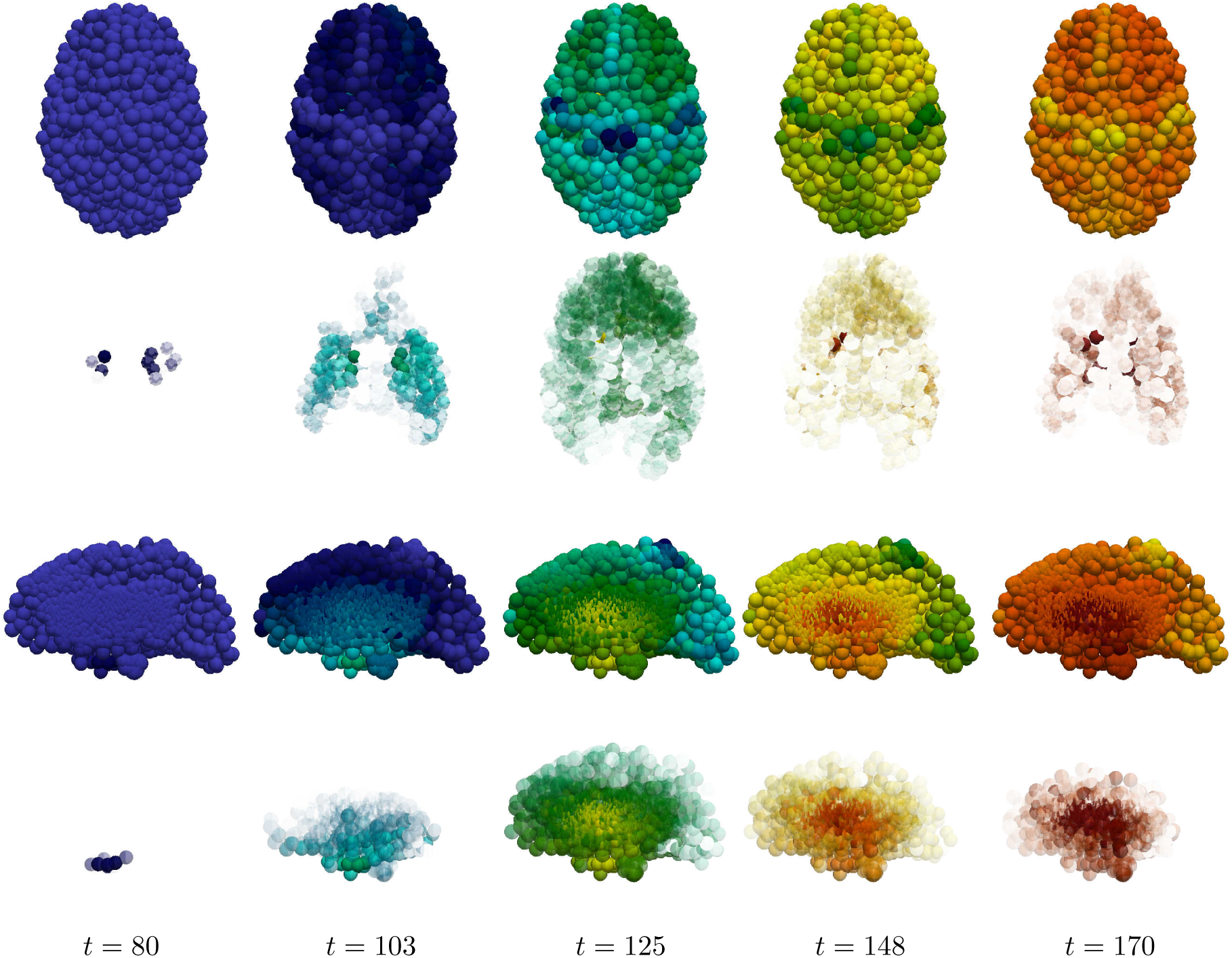
Damage progression in primary tauopathy. Horizontal plane view (top row) with opacity exaggerated (second row) progression. sagittal view (third row) with opacity exaggerated (fourth row) progression. Dark blue indicates the minimal damage value of *q* = 0.0; bright red indicates the maximum of *q* = 1.0. Intermediate values are: purple (*q* = 0.14), sky blue (*q* = 0.29), green (*q* = 0.43), yellow (*q* = 0.57), orange (*q* = 0.71), and dark red (*q* = 0.86).

**Figure 24:**
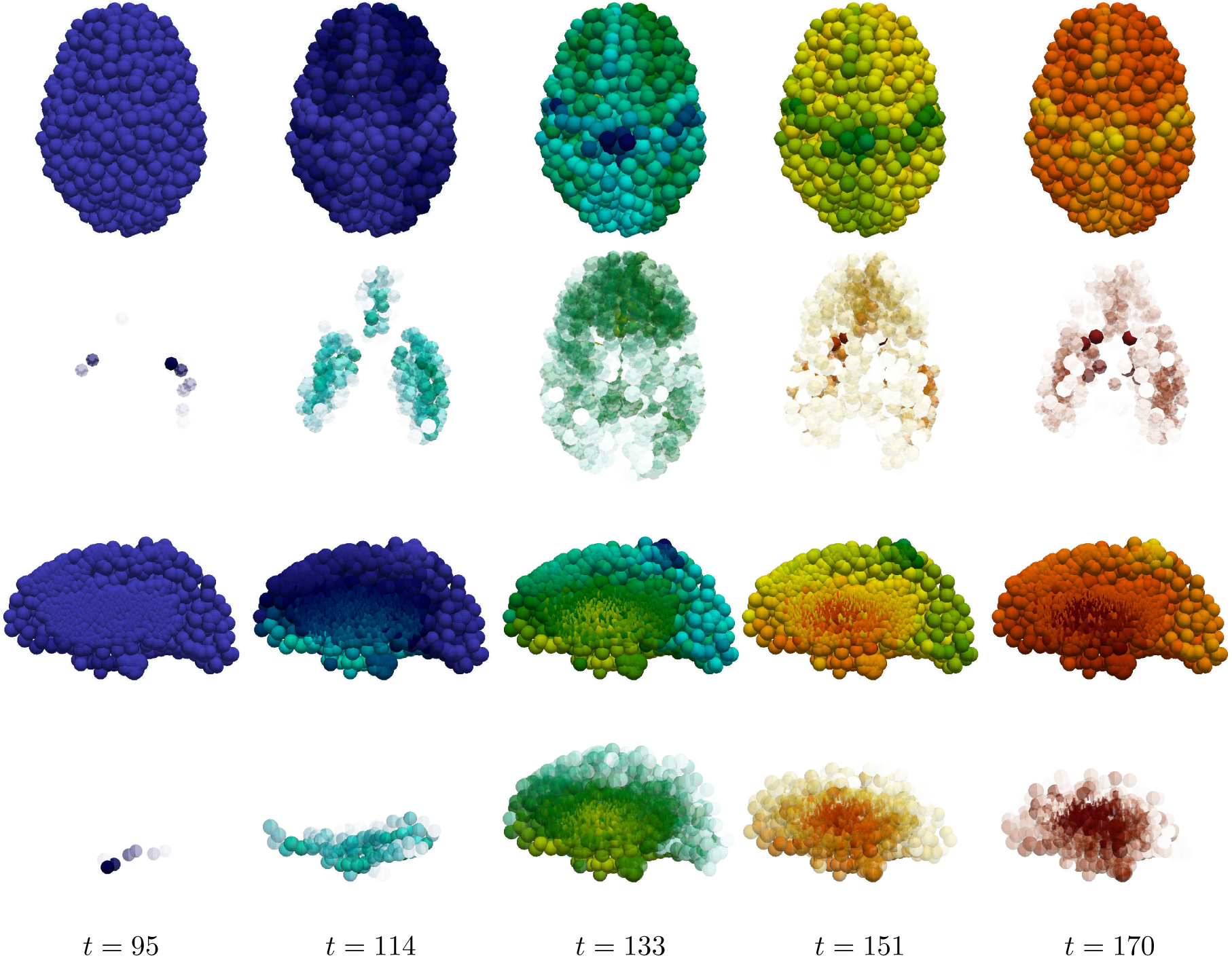
Damage progression in secondary tauopathy. Horizontal plane view (top row) with opacity exaggerated (second row) progression. sagittal view (third row) with opacity exaggerated (fourth row) progression. The color scale is identical to that of Figure 23.

It is challenging to discern differences between the fully opaque horizontal views of Figure 23 v.s. Figure 24; some discrepancies are apparent in the sagittal views, however. Relative opacity exaggeration is used to gain further insight. At each time the minimum and maximum damage, denoted *D*_min_ and *D*_max_, was computed across all regional nodes of the brain connectome; opacity was then set to linearly increase from: fully transparent at the average 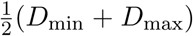; to fully opaque at the maximum value *D*_max_. The resulting opacity exaggeration scheme shows, at each time step, the relative distribution of the most damaged regions.

The aforementioned opacity scheme leads to a further observations. First, the distribution of relative significant damage in primary tauopathy (Figure 23, second and fourth rows) is clustered more centrally to the toxic *τ* P seeding site of the transentorhinal cortex. Conversely, the distribution of relative significant damage in secondary tauopathy (Figure 24, second and fourth rows) is distributed in the direction of the temporobasal region; a site associated with A*β* seeding. As the disease progresses, *t* = 103 and *t* = 114 in Figures 23 and 24 respectively, we see two distinct differences: relative damage is more connected, in the horizontal plane, in addition to more diffuse in the coronal direction, of the sagittal plane, for the case of primary tauopathy; in secondary tauopathy the relative damage in the horizontal plane forms three distinct clusters while severe damage in the sagittal plane is follows the temporobasal and frontomedial directions.

It is increasingly difficult to visually detect qualitative patterns in later stages of significant damage progression; that is, *t ≥* 125 for primary tauopathy and *t ≥* 133 for secondary. Nevertheless it appears that late stages, *t* = 148 and *t* = 170, for primary tauopathy display a more diffuse distribution of significant relative damage away from the transentorhinal region; whereas late secondary tauopathy, *t* = 151 and *t* = 170, show more comparative significant damage in the areas associated with A*β* initial seeding.

Taken collectively: these observations suggest that damage onset and the relative distribution of severe damage may offer distinct points of view for application modelling to both typical Alzheimer’s disease along with its neuropathological subtypes [77, 78].

## 6 Further observations and concluding remarks

In this section we reflect on the analytic and computational results of the manuscript. In Section 6.1 we list some advantages and disadvantages of the current perspective; in Section 6.2 we endeavor and discussion of several aspects of the work and avenues of open, or further, enquiry; and in Section 6.3 we offer brief concluding remarks. The proposed model is based on physical protein aggregation kinetics; the simplest such two-famiy-two-species interacting protein model one could posit. Nevertheless, the model is mathematically sophisticated enough to evince two distinct pathology regimes, termed primary and secondary tauopathy, of potential clinical interest. After discretizing (1) on a structural connectome: an approachable system of non-linear ordinary differential equations, (6)-(9), emerges which can be solved using standard mathematical software; such as Mathematica or Matlab. As a result, we expect the model to be widely appealing to the computational neurodegenerative disease community as a starting point for gaining further insight into protein-protein interactions in the context of Alzheimer’s disease.

### 6.1 Advantages and limitations

The deterministic nature of the model, c.f. (1), has at least three distinct advantages over complex, stochastic models: first, the reaction terms of (1) represent simplified [49, 50], but physical, protein aggregation kinetics with a basis in experimental measurement [50, 52, 53, 54, 56]; second, the connectome-discretized equations, (6)-(9), can also be readily implemented using off-the-shelf mathematical software (e.g. Mathematica or Matlab, etc). Thus, (6)-(9) are easily approachable and do not require probabilistic postulations, based on data or otherwise, regarding underlying distributions. A third advantage is that (6)-(9) are amenable to an a-priori mathematical analysis. This analysis is immutable in nature and much can be observed (c.f. Section 6.2) as a result of using standard methods from the theory of ordinary differential equations and non-linear diffusion-reaction systems. Conversely, probabilistic models may need extensive tuning, reformulation or data curation in order to determine a model’s emergent properties. An independent investigation, i.e. model fitting and application, founded on datasets with differing fidelity may produce divergent results. Such models essentially act in service to deeply mine a set of data but are not always directly helpful to elucidate the impact of individual disease mechanisms.

Conversely, (6)-(9) has inherent limitations. As discussed in recent literature: [35, 79] there are challenges surrounding the acquisition of the parameters in deterministic models such as (6)-(9). In vitro kinetic parameters, regulating the multiplication and growth of several proteins, have been ascertained for A*β, τ* P, *α*-synuclein and others; c.f. the citations in [79]. If we disregard the clearance terms in the prototypical heterodimer model, (6)-(7), then precisely two kinetic coefficients remain: source production (*a*_0_) and healthy-to-toxic conversion (*a*_2_). The in-vitro experimental estimation of protein-specific aggregation kinetic parameters, however, typically relies on more complex theoretical models: consisting of at least five kinetic parameters; and an infinite number of equations (c.f. for instance [48, Sec. 3.2]). It is therefore not immediately clear how to obtain explicit values for the kinetic rate parameters of (6)-(9). Asymptotic expansions have provided links between the rate coefficients of other more complex models and their simpler counterparts, c.f. [Sec. 2.2][47], and such approaches may provide insight into reducing experimental parameters from the five-parameter models [79, 48] to those of (6)-(9). A further complication, though, is that even if explicit, experimentally verified, in-vitro parameters were available for (6)-(9) these do not necessarily translate into the correct parameters in vivo [79] where indirect, and locally varying, mechanisms (such as aggregates interacting with cells, or the effects of inflammation on aggregation dynamics) may play a role in altering the associated rates; either globally or regionally.

### 6.2 Discussion and open questions

Neurodegenerative diseases are complex and multi-scale processes. The point of view of (6)-(9) is to reduce this complexity by considering a collection of aggregate mechanisms and their implications. For instance (1) can be viewed, more conversationally, as the following collection of general mechanisms: (a) there exists two protein families; (b) each family has a healthy and toxic species; (c) these species are produced and cleared at some aggregate (regional) rate (d) any movement of these species, within the brain, is primarily determined by the macroscale axonal structure; (d) healthy proteins within a family can become toxic, at some (regional) rate, based on the presence of other toxic proteins of that family; and (e) the conversion of healthy-to-toxic proteins, for the second family, is further influenced by the presence of the toxic population of the first family. The current literature suggests that this collection of observations outlines a minimal prion-like model of Alzheimer’s disease progression.

In section 6.1 it was mentioned that one advantage of simple deterministic models, such as (6)-(9), is that the impact of individual mechanisms can be elucidated and several emergent behaviors can be ascertained a priori. Models such as (1) therefore lead, naturally, to additional questions and serve as a trailhead for further development. The first, and critical, observation is that: (6)-(9) implies that the local balance of clearance, e.g. (21), plays a *fundamental* role in disease initiation. In light of the seminal work of Braak and Braak [11] this leads naturally to the question: what are the local (toxic *τ* P) clearance properties characterizing the transentorhinal region, (which defines the early Braak stages) and how do these local properties differ from other regions? Aspects of the fine-scale clearance mechanisms of toxic *τ* P remain unclear or are even controversial [80, 81, 82, 83]. Nevertheless, our simple framework reinforces the sentiment echoed by experimentalists: that understanding these processes may be critical to a mechanistic understanding of the initiation of the disease cascade.

A second observation emerging from (6)-(9) is that the progression of Alzheimer’s disease may consist of a confluence of brain regions simultaneously in differing states characterized by contrasting fundamental dynamics. In particular, section 3.4 shows that even a simple model of AD development suggests potentially complex disease phenomenology; one where *τ* P can evolve independently of A*β* (termed primary tauopathy) and one where *τ* P depends intrinsically on the presence of A*β*. Furthermore, the line between these two regimes is demarcated by: the balance of local clearance; and the degree of local influence of A*β* on the toxification of *τ* P [12, 26, 30, 31, 32, 33] as expressed by the bulk parameter *b*_3_. Depending on these local attributes we could have some areas of the brain in a state of primary tauopathy and others in a state of secondary tauopathy; the latter regions having their tauopathy delayed until a toxic A*β* population is established while the former regions are free to develop toxic *τ* P and NFT independently. This leads naturally to another fundamental line of further enquiry: what are the simplest additional relations, extended (6)-(9), needed to suitably describe the evolution of clearance and toxicity rates alongside protein pathology?

Our simple mathematical model suggests, as a third observation, that the rate of toxic A*β*-*τ* P interaction (i.e. *b*_3_) is not a passive facet of disease phenomenology but, rather, may play a much more integral role. We have already discussed, above, that *b*_3_ plays a role in secondary tauopathy; it can do this by lowering *v*_4_, in (15), thus ensuring that 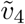 is an admissible state. Other interesting observations regarding *b*_3_ were discussed in sections 5.1.1, 5.1.2 and 5.2. The observation in 5.2 is straightforward so we will mention it first: namely, that transneuronal damage propagation in the model has a ‘minimum speed’ in both primary and secondary tauopathy; and that increasing *b*_3_ has no effect. In particular Fig. 22, and the discussion in Section 5.2, suggest that lowering the ‘transmission coefficient’, reflected in *k*_4_, below a certain threshold does not lower transneuronal damage; and that increasing *b*_3_, for a fixed choice of *k*_4_, does not increase the overall propagation of damage. These details lead to a somewhat interesting observation: that the rate of neuronal damage from the structural *network topology* of the brain may exhibit a *baseline*, or minimum, value; independent of the details driving local damage (e.g. from local toxicity).

In sections 5.1.1 and 5.1.2 we discussed two observed features of tauopathy directly related to *b*_3_: the time of onset and the *τ* P ‘invasion window’. We recall that: the time of onset is defined as the first appearance of toxic *τ* P while the ‘invasion window’ is the timespan starting at a 1% toxic *τ* P concentration and terminating when the asymptotic steady-state value is achieved. In primary tauopathy (section 5.1.1): disease onset time is virtually unaffected by varying *b*_3_ whereas increasing levels of *b*_3_ shortens the tauopathy invasion window. In addition, the asymptotic concentration value of toxic *τ* P increases with *b*_3_ so that, overall, increasing *b*_3_ implies that a more severe tauopathy will develop, faster, at a similar starting point in time (c.f. Figure 14). The picture in secondary tauopathy (section 5.1.2) is different. We see, again, that increasing *b*_3_ does increase the severity of the tauopathy (Figure 16b); however, this is where the similarities with primary tauopathy end. First, as *b*_3_ increases the time of onset decreases (Figure 16b). Second, the invasion window in secondary tauopathy does not decrease monotonically with decreasing *b*_3_ (Figure 16c); rather, we see the invasion window start time and end both decrease, with increasing *b*_3_, while the start time decay and end time decay, relative to increasing *b*_3_, is different (Figure 17). This is the cause of the initial drop, from *b*_3_ = 0.75 to *b*_3_ = 3.0, of the invasion window in Figure 16c followed by an increase to a steady invasion window length circa *b*_3_ = 12. The observation that increased *b*_3_ can decrease the time of onset in secondary tauopathy, which requires the presence of A*β*, while also impacting the invasion window time is reminiscent of the effects associated to the presence of particular Apoliprotein E (APOE) allele configurations. For instance: APOE *E*4 carriers are more likely to develop AD; toxic A*β* production and deposition is more abundant in APOE *E*4 carriers; and APOE *E*4 exacerbates A*β*-related neurotoxicity [84].

In section 5.1.3 we discussed a mixed-modality instantiation of the model (6)-(9), with some regions in a state of primary tauopathy with all others in a state of secondary tauopathy, using hand-selected synthetic parameters. It was demonstrated that such a mixture of states can reproduce salient features seen in Alzheimer’s disease; in particular, the distinct distribution patterns [68, 69] of 18F-AV-1451 radiotracer are clearly observed. This observation suggests that the model of (6)-(9) is sufficiently rich and implies that the undertaking of a comprehensive data fitting and comparison study is both well warranted and an optimistic endeavor. It is interesting to note, though, that the distinction between primary and secondary tauopathy is not simply one of differently-valued parameters; in particular the two states are differentiated by the balance of clearance inequalities discussed in Sec. 3.4. In particular, we have *ã*_1_*/a*_2_ *< a*_0_*/a*_1_ and 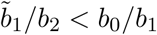 for primary tauopathy; for secondary tauopathy the latter inequality changes sign to 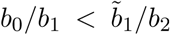. This observation implies that an arbitrary parameter fitting could produce accurate results, compared to data, while still being questionable since the fitting would imply secondary characteristics regarding regional clearance attributes which may or may not hold. It would therefore be beneficial to carefully consider a data-based measure of regional clearance, for both A*β* and *τ* P, when selecting a data fitting method; possibly incorporated as a constraint or as part of a cost functional. Nevertheless, the dual regimes of primary and secondary tauopathy provide a verdant backdrop for further modelling endeavors; both in terms of fitting to clinical imaging data and to probing and modelling possible ties between *b*_3_, APOE configuration, and secondary tauopathy.

In section 1 we mentioned that the role of Tau in AD formation, and development, is beginning to be recognized as a potentially significant factor. Despite this, open questions about the nature of tau, and tauopathy, in AD remain. For instance: one could argue that healthy *τ* P, being bound to microtubules, should not be diffusing at all. The literature suggests that even healthy tau in healthy neurons exhibit mobility within the cell [85], is secreted into the extracellular space [86, 87], and that extracellular tau is taken up by neighboring neurons [88]; even in the absence of pathology. It is not entirely clear what the correct choice of *ρ*, in (5), should then be. Despite the literature seeming to suggest that *ρ >* 0 one could still insist that *ρ* ≪ 1, or possibly even assert that *ρ* = 0, should be chosen in (5) for the graph Laplacian of (8). Regarding impacts to the model: this perspective alone would not affect any of the analytic observations of Section 3. Indeed, if *v* is the vector whose *j*^th^ entry is the healthy tau concentration, *v*_*j*_, in node *j* then if *v* is constant, or nearly so, then the graph Laplacian applied to *v* is zero, or nearly so, regardless of the value of *ρ*. Thus, since all of the nodes in the computational investigations of Section 4 had their healthy tau populations set to the same constant value: the effect of any healthy tau diffusion in the simulation results there would be expected to be entirely negligible as well.

The nature of the rates for healthy tau production and clearance, in the literature, are also not fully understood. Indeed, the visual confirmation of the mRNA machinery for localized transcription [89] of tau in axons, and growth cones, is less than two decades old; clarifying important aspects of tau clearance, both healthy and diseased, is an ongoing process [80, 82, 83]. Our results indicate that violation of the balance of clearance inequalities, (21), is fundamental for disease initiation and phenomenology; for instance: if healthy tau were not regenerated, so that *b*_0_ = 0, then the regime of ‘primary tauopathy’ (which requires that 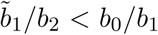) would be an impossibility. This would imply that, in the context of our model, that the development of all tauopathies would require an accompanying amyloidopathy and would seem to preclude those tauopathies which are mostly dominated by toxic *τ* P spreading [8]. It has been observed that: tau is expelled from neurons [86], including healthy ones, on a periodic basis [87]; and that tau plays a role in cell signalling, cell polarity, synaptic plasticity and the regulation of genomic stability [90, 91]. These observations, alongside recent work in adult neurogenesis [92], give good reason to suspect that both *b*_0_ *>* 0 and *b*_1_ *>* 0; at least in the healthy brain and early in disease progression. An open question, though, is how these quantities may change with disease progression. For instance, one could extend the current model by coupling *b*_0_ with the damage coefficient *q*; reflecting the fact decreased healthy tau synthesis could result from neuronal loss, and the decline of neurogenesis [92], throughout AD progression.

The final observation we mention regards an open question surrounding the imaging, and construction, of the structural connectomes used in such network models. We have used an often cited connectome [58, 59]. This connectome is available in various resolutions; the lowest of which consists of 83 vertices (regions of interest) while the highest resolution case, which we have used here, consists of 1015 vertices. However, there are apparent differences in both A*β* and *τ* P staging when solving equations, such as (6)-(9), on the low versus high resolution connectomes. We used the simple, illustrative parameters described for primary tauopathy, in Section 5.1.1, and recorded the average regional tau concentration at six fixed time points; the time points were selected to span disease progression. Figure 25 shows the results of this tau staging experiment for nine regions. We can see that the two different resolutions of connectomes, derived from the same set of patient data, offer distinct staging patterns for tau progression. This implies that the connectome itself may play a significant role in retrieving results that match clinical data. Since the parameters of (6)-(10) have a physiological interpretation a simple ‘fitting’ to available clinical data is not satisfactory. Figure 25 suggests that developing a more rigorous understanding of computational staging behavior should be endeavored seriously and from first principles. Validating computational (tau) staging behavior, at different connectome resolutions, against clinical standardized-uptake-value-ratio (SUVR) studies, e.g. [93], is an important next step.

**Figure 25:**
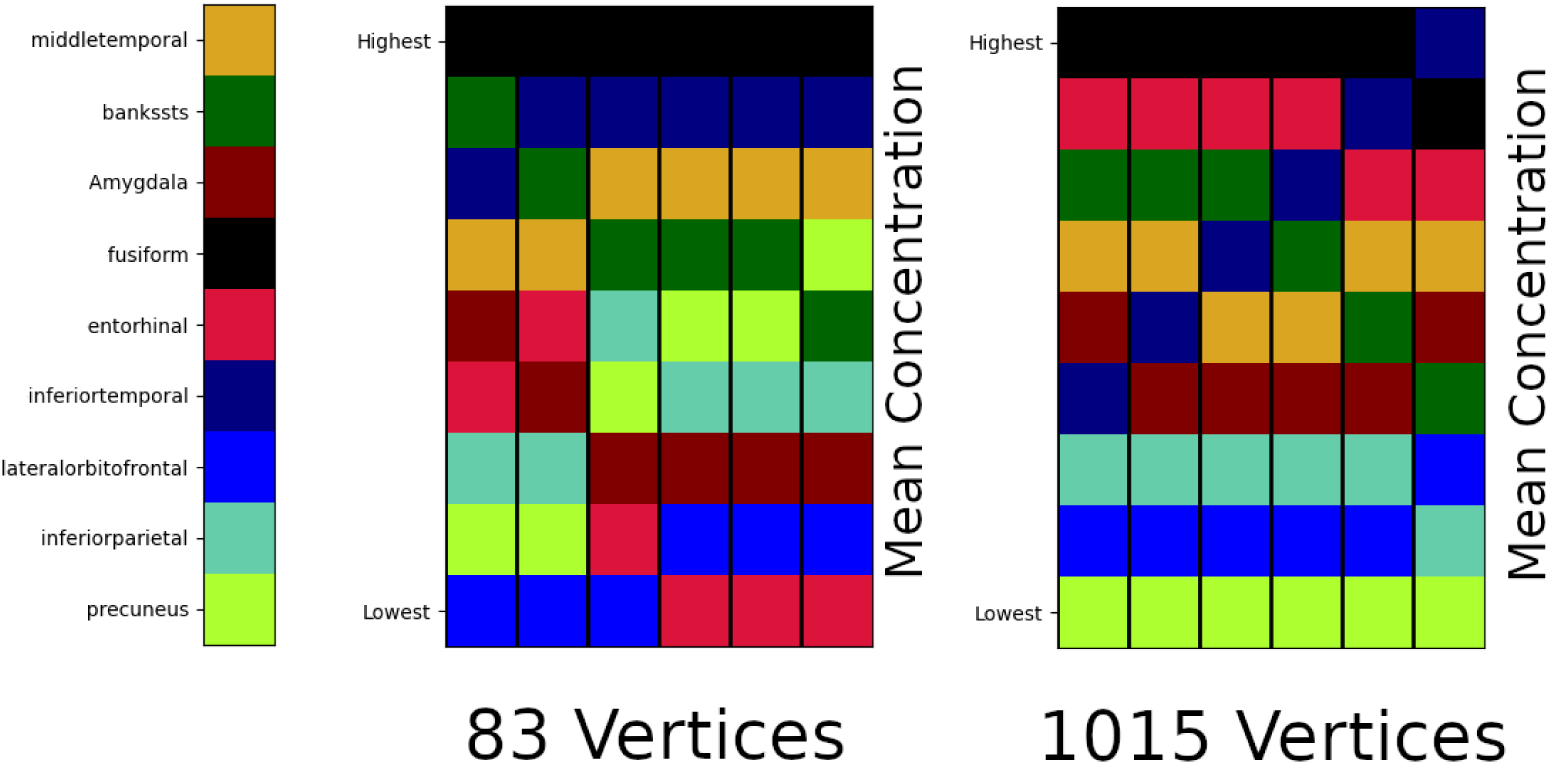
Toxic *τ* P average regional concentration; six fixed time points. 83 (left) versus 1015 (right) vertex connectomes

### 6.3 Concluding remarks

We have presented a novel, minimal, and deterministic theoretical mathematical model of protein propagation that includes two interacting protein species. The model is motivated by recent experimental evidence regarding the potential importance that interactions between A*β* and *τ* P may play in the development of AD pathology [14, 16, 23, 24, 25]. The primary contributions of the current manuscript are: clearly, and mathematically, establishing the intrinsic dependence of the model on the balance of clearance inequality, (21), in Section 3.4; the stability analysis of the modes of primary and secondary tauopathy in Sections 3.3 and 3.4; and establishing the speed of propagation of toxic fronts in Section 3.5. Further novel contributions of interest include: demonstrating qualitative properties of disease propagation and damage, in primary and secondary tauopathy (Section 4), using globally constant, but non-physical, parameters. In particular, we have seen that the topology of the brain connectome leads to complex behavior in both pathological regimes. Finally, in Section 6.2 we have contextualized numerous analytic and computational observations with reference to the current literature and drawn attention to open avenues of further research suggested by the current work.

Alzheimer’s disease is a complex and multi-scale disease. The need for mathematical models, presenting observed disease characteristics, that are computationally tractable is pressing. Our findings suggest that further enquiry into both protein interaction and clearance processes is an important path forward in elucidating key mechanisms in the progression of these diseases. Due to the ease of implementation of (6)-(10), and the widespread interest in computational neurodegenerative disease, we hope that this model will be appealing, to the community, for probing the nuances of protein-protein interactions in neurodegenerative disease development.

## Supporting information

Supplemental movies (all, compressed)

## Acknowledgments

Data used in the preparation of this article were obtained from the Alzheimer’s Disease Neuroimaging Initiative (ADNI) database (adni.loni.usc.edu). The ADNI was launched in 2003 as a public-private partnership, led by Principal Investigator Michael W. Weiner, MD. The primary goal of ADNI has been to test whether serial magnetic resonance imaging (MRI), positron emission tomography (PET), other biological markers, and clinical and neuropsychological assessment can be combined to measure the progression of mild cognitive impairment (MCI) and early Alzheimer’s disease (AD). For up-to-date information, see www.adni-info.org.

This work was supported by the Engineering and Physical Sciences Research Council grant EP/R020205/1 to Alain Goriely, by the National Science Foundation grant CMMI 1727268 to Ellen Kuhl, and by the John Fell Oxford University Press Research Fund grant 000872 to Travis Thompson.

## A Numerical verification

In this appendix we test our computational platform by recovering the basic homogeneous dynamics of the full network model. To do this we use two hypothetical sets of illustrative, non-clinical parameters; one set of parameters for each regime. In Section A.1 we illustrate the four possible patient states (stationary points) of Section 3.2. In Section A.2 the primary and secondary tauopathy, c.f. Section 3.3, patient state transitions are simulated and model patient dynamics are discussed in more detail. Front propagation in the brain connectome network is confirmed using synthetic left-right hemisphere initial seedings in Section 4.1.

### A.1 Patient states of the network system

We now briefly illustrate the four stationary states of the homogeneous system discussed in Section 3.1. To demonstrate that each of the predicted stationary points is indeed a stationary point of the homogeneous network system, c.Section 3.1, we select illustrative parameters that satisfy the requisite characterizing inequalities. Every node in the brain network is then seeded with the initial value corresponding to the selected fixed point. We expect, and demonstrate, that the system remains stable at that fixed point.

We will confirm the stationary points by selecting the effective diffusion constant, *ρ* of (5), as unity and solving (6)-(10) for *t* ∈ [0, 10] using one thousand time-steps. For the healthy A*β*-healthy *τ* P state, c.f. (12), we select *a*_0_ = 0.75 and *b*_0_ = 0.5; all other parameters are set to unity. All nodes were seeded with the corresponding initial value

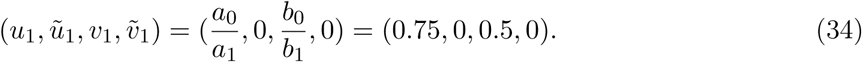

Figure 26a shows the plot of global mean tracer concentration with time and confirms that the healthy A*β*-healthy *τ* P state is stationary under the given conditions. For the healthy *τ* P-toxic A*β* fixed point, c.f. (13), we begin with the previous parameters and reduce the toxic A*β* clearance by 40%. We therefore have *ã*_1_ = 0.6 and keep the previous parameters fixed. We then have

**Figure 26:**
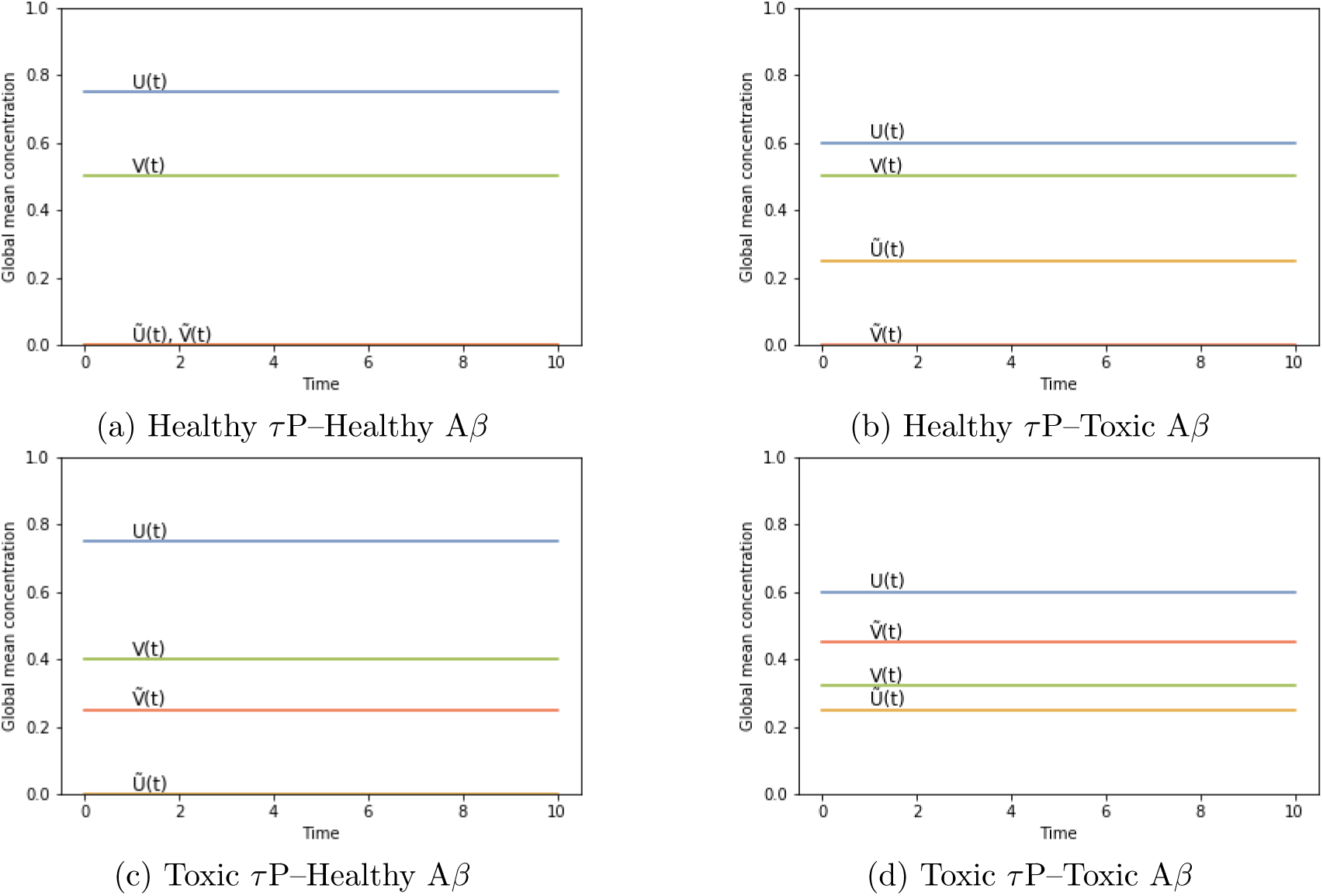
Computational verification of the stationary points (12)-(15)

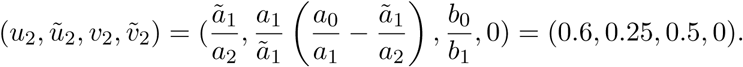

The stationary behavior is again demonstrated; c.f. Figure 26b. For the third stationary state, given by (14), we begin once more with the parameters of the healthy A*β*-healthy *τ* P state and reduce the toxic tau clearance parameter by 60%. We then have 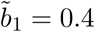 and keep all other parameters as in the healthy A*β*–healthy *τ* P state. All nodes are then set to the corresponding initial value

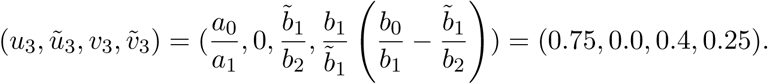

Once more, Figure 26c, we see the stationary characteristic we expect. For the final stationary point, c.f. (17), we use the reduced toxic clearance parameters from the second and third stationary points above, *ã*_1_ = 0.6 and 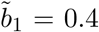, in addition to the original production values, *a*_0_ = 0.75 and *b*_0_ = 0.5, of A*β* and *τ* P respectively. All other parameters not explicitly mentioned are again taken to be unity. Given these choices we can directly compute *µ* and *v*_4_, via (16)-(17), as

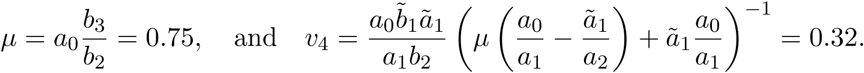

Using the above, along with the expressions for *v*_1_, *v*_3_, *u*_1_, *u*_2_ and *ũ*_2_ from (12)-(14), the value of 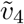 is given directly from the fourth entry of (17) as

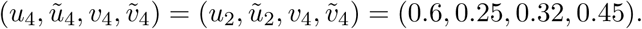

The final plot, for the fourth stationary point, is shown in Figure 26d. Coronal and sagittal plane views of the stationary point verification computation at *t* = 10 are shown in Figure 27.

**Figure 27:**
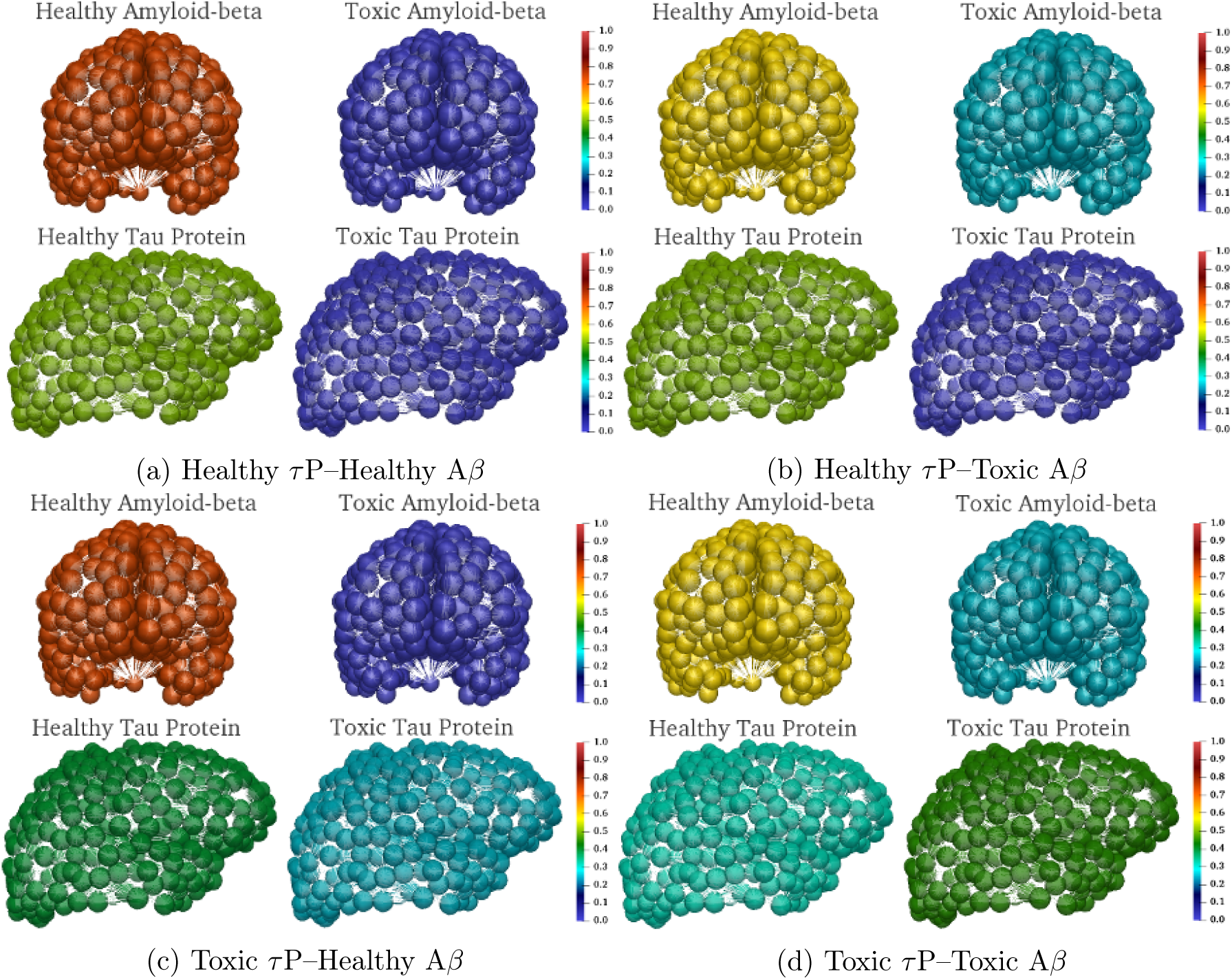
Network system stationary point realization; coronal (top) and sagittal (bottom) views.

### A.2 Patient pathology transitions of the network system

We briefly illustrate the homogeneous state dynamics of the network system; verifying the theoretical view of Section 3.3 on the complex brain network geometry of Figure 2.

#### A.2.1 Primary tauopathy

We consider a hypothetical susceptible model patient characterized by the parameters chosen in Appendix A.1. All four of the stationary points discussed in Section 3.2 coexist with this choice of parameters; hence, these parameters fall into the regime of primary tauopathy. In this section we verify the homogeneous state transitions, between the states of Figure 27, of (6)-(10) discretized on the brain network geometry of Figure 2. The selected illustrative primary tauopathy parameters are collected in Table 5 for posterity.

The eigenvalues, (19) and (20), at the healthy A*β*–healthy *τ* P stationary point (*u, ũ, v*, 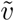) = (0.75, 0, 0.5, 0) can be calculated. We see that *λ*_*Aβ*,1_, *λ*_*τP*,1_ *<* 0, i.e. stable to healthy A*β* and *τ* P perturbations, while *λ*_*Aβ*,2_, *λ*_*τP*,2_ *>* 0 so that the otherwise healthy patient brain is susceptible to perturbations in both toxic A*β* and toxic *τ* P. Utilizing the given parameters to evaluate the stability properties at the second stationary point, (*u, ũ, v*, 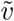) = (0.6, 0.25, 0.5, 0) c.f. (13), we have *λ*_*Aβ*,1_, *λ*_*Aβ*,2_, *λ*_*τP*,1_ *<* 0 and *λ*_*τP*,2_ *>* 0; at this state the patient is susceptible only to a perturbation in toxic tau. Likewise at the third stationary point, (*u, ũ, v*, 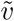) = (0.75, 0, 0.4, 0.25) c.f. (14), we have *λ*_*Aβ*,1_, *λ*_*τP*,1_, *λ*_*τP*,2_ *<* 0 and *λ*_*Aβ*,2_ *>* 0 so that the patient in this state is only susceptible to an addition of toxic A*β*. Finally the fixed point (15) is fully stable, i.e. all eigenvalues are negative, and no further disease transition is possible from this state.

Verifications of the primary tauopathy homogeneous state transitions, first depicted in Figure 3, for the full connectome simulation are shown in Figure 28. For instance the healthy state, (*u*_1_, *ũ*_1_, *v*_1_, 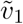), perturbation with respect to both toxic A*β* and toxic *τ* P results in the fully toxic state, (*u*_4_, *ũ*_4_, *v*_4_, 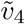); this is shown in Figure 28c and appears in Figure 3 as the blue (diagonal) path.

**Figure 28:**
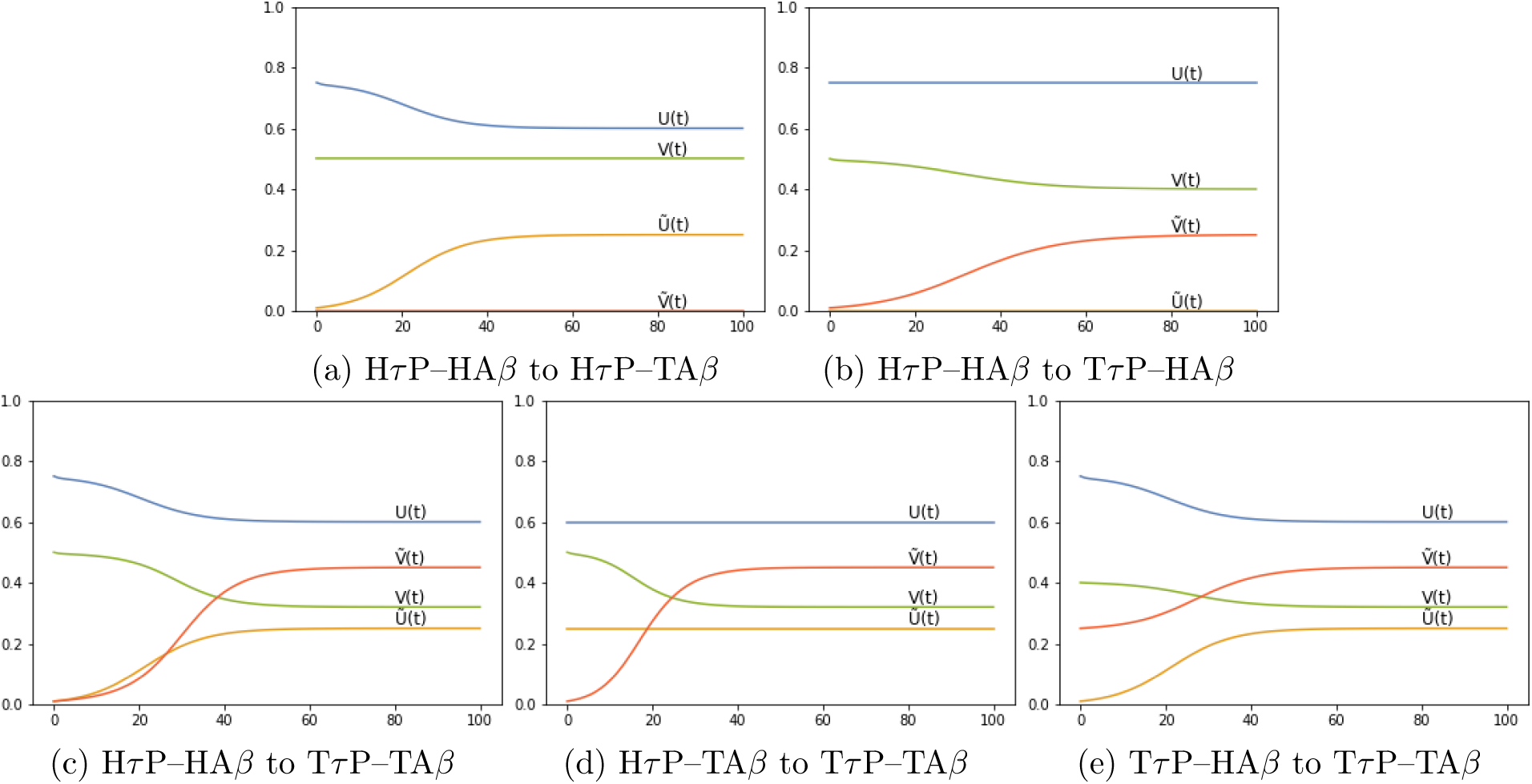
State transitions in primary tauopathy. Concentration (y axis) vs. simulation time.

#### A.2.2 Secondary tauopathy

The secondary tauopathy disease model arises when *v*_1_ *< v*_3_, so that the stationary point (14) is in an unphysical state, while (12), (13) and (15) remain well defined. One way that this can be achieved is for *b*_3_, the coefficient mediating the effect of toxic A*β* protein on inducing healthy tau toxification, to be such that both *v*_4_ *< v*_1_ and *v*_4_ *< v*_3_; a decrease in *b*_2_ can also accomplish this goal, c.f. (15).

The condition *v*_1_ *< v*_3_ is equivalent to 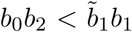. One can transform the primary tauopathy patient described by the parameters of Table 5 to a secondary tauopathy patient by decreasing *b*_2_ by twenty-five percent; from 1.0 to 0.75. In this regime *v*_1_ = 0.5 and 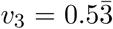 and the stationary point (14) is physically inadmissible. The admissible stationary states are

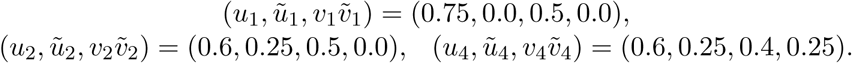

We see that the first and second stationary points are identical to the case of primary tauopathy and the fourth is perturbed in the (*v*, 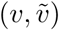) components. Strictly speaking, the healthy patient in this regime is susceptible only to toxic A*β* infection; that is *λ*_*Aβ*,1_, *λ*_*τP*,1_, *λ*_*τP*,2_ *<* 0 and *λ*_*Aβ*,2_ *>* 0 at (*u*_1_, *ũ*_1_, *v*_1_, 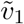). Verification of the healthy state robustness to perturbations in toxic tau, 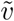, is shown in Figure 29.

**Figure 29:**
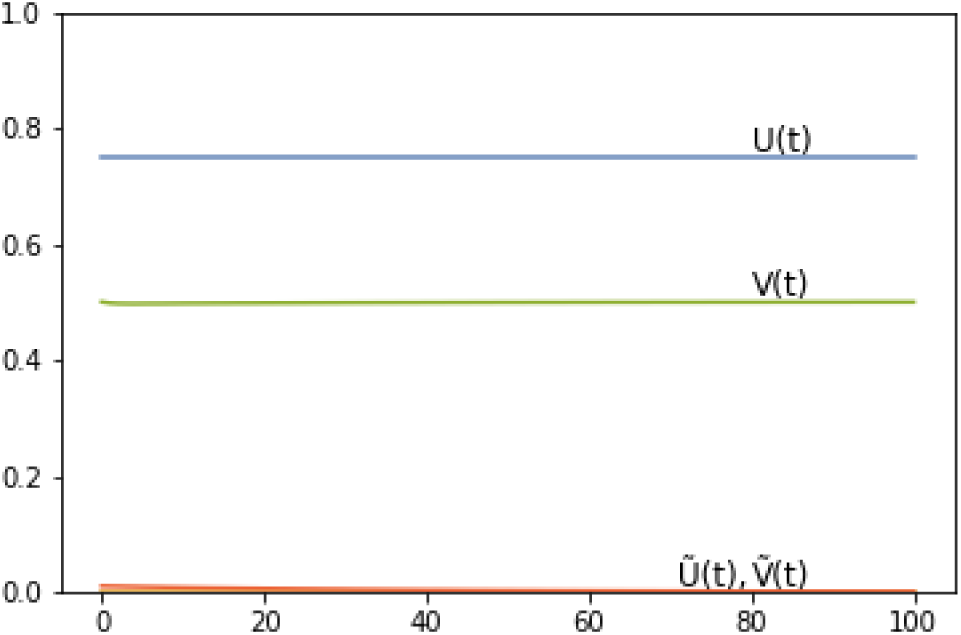
H *τ*P-HAβ,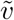 stable

At the healthy state *λ*_*Aβ*,2_ *>* 0 holds. Thus, the susceptible, but otherwise healthy, secondary tauopathy patient is at risk of directly developing A*β* proteopathy. This is verified by perturbing the healthy state by a small concentration in *ũ*; the pursuant transition from the Healthy *τ* P– Healthy A*β* state to the Healthy *τ* P–Toxic A*β* state is pictured in Figure 30b. Having arrived at (*u*_2_, *ũ*_2_, *v*_2_, 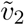) the patient is now susceptible to tauopathy as *λ*_*τP*,2_ *>* 0 there; perturbing 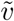 then develops to the Toxic *τ* P–Toxic A*β* state as shown in Figure 30c.

**Figure 30:**
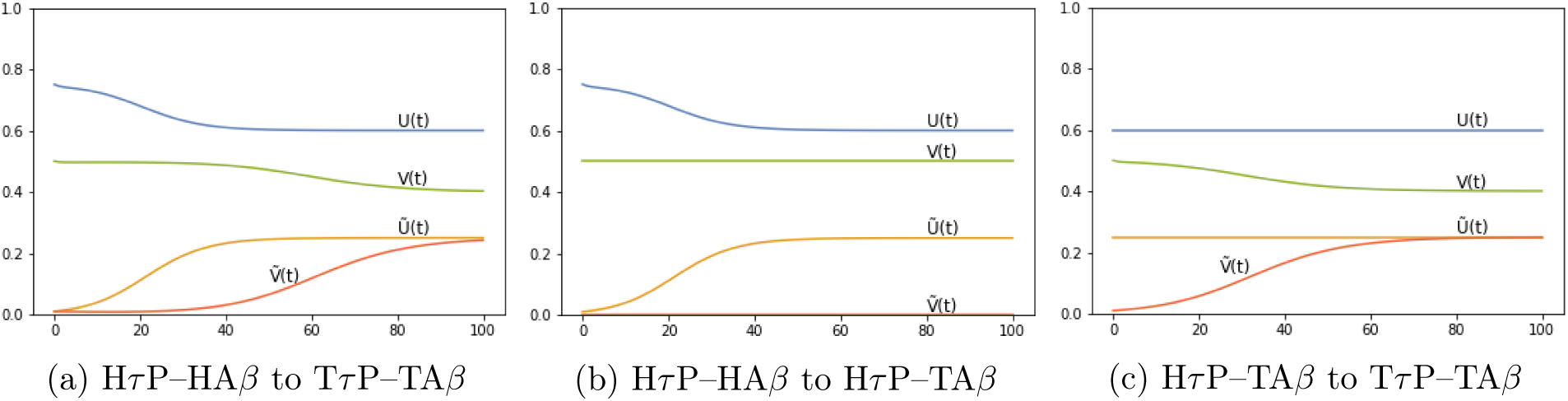
State transitions in secondary tauopathy. Concentration vs. simulation time.

In fact, as postulated in Section 3.3 c.f. Figure 4, the fully diseased state (*u*_4_, *ũ*_4_, *v*_4_, 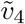) is reachable from the healthy state provided that toxic A*β* is present alongside some toxic tau perturbation. This can be seen directly from *λ*_*τP*,2_ in (20). Consider the Taylor expansion of (20), evaluated with *b*_2_ = 0.75 and all other parameters as in Table 5, about 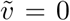. We first set *θ* = *ũ* + 0.6 and we let 0 *≤ ϵ* ≪ 1 be denote a small perturbation in 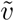. It is evident that the effect on *λ*_*τP*,2_ due to a perturbation in toxic tau depends here on both toxic amyloid, *ũ*, and healthy tau, *v*, concentration levels. Then, using that *ũ ≥* 0, and *v ≥* 0, we approximate (20), to order *ϵ*^2^, around 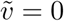 by

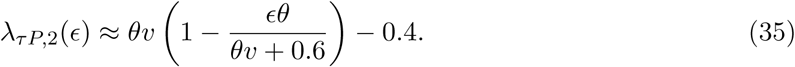

If we presume, for instance, that the susceptible secondary tauopathy patient has healthy levels of tau protein, i.e. that *v* = *v*_1_ = 0.5, we can directly visualize the effect of toxic A*β* on *λ*_*τP*,2_. Figure 31 shows the approximate value of *λ*_*τP*,2_ (y-axis, c.f. (35)) versus the toxic A*β* value *θ*(*ũ*) = *ũ*+ 0.75 (x-axis) for three given perturbations *E*. Evidently, as *E* decreases the effect of *ũ* on increasing *λ*_*τP*,2_ is not diminished. Thus an initial toxic *τ* P seed will develop into a full blown infection provided *ũ* is present, or quickly develops, in sufficient quantity to evolve *λ*_*τP*,2_ above zero. This is precisely the behavior predicted iSection 3.3 (Figure 4). In accordance we see, c.f. Figure 30a, that perturbing both *ũ* and 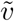 simultaneously from the initial healthy state induces direct evolution to fully diseased state.

**Figure 31:**
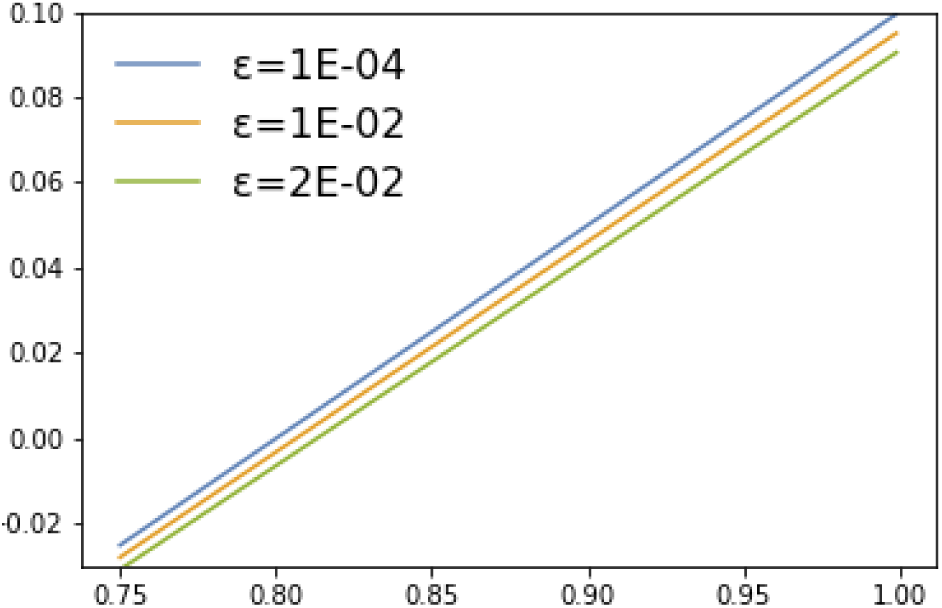
*λ*_*τP*,2_ vs. *θ* = *θ*(*ũ*)

## B Supplementary Data

S1 Movie: Front dynamics in primary tauopathy: visualization of an illustrative model simulation using a synthetic channel domain.

S2 Movie: Front dynamics in primary tauopathy: visualization of an illustrative model simulation using a high-resolution structural brain connectome domain.

S3 Movie: Front dynamics in secondary tauopathy: visualization of an illustrative model simulation using a synthetic channel domain.

S4 Movie: Front dynamics in secondary tauopathy: visualization of an illustrative model simulation using a high-resolution structural brain connectome domain.

S5 Movie: Toxic A*β* in primary and secondary tauopathy: visualization of the toxic A*β* species in an illustrative model simulation using a high-resolution structural brain connectome domain. Qualitative propagation of the A*β* species was identical in both primary and secondary tauopathy.

S6 Movie: Toxic *τ* P in primary tauopathy: visualization of the toxic *τ* P species in an illustrative model simulation of primary tauopathy using a high-resolution structural brain connectome domain.

S7 Movie: Toxic *τ* P in secondary tauopathy: visualization of the toxic *τ* P species in an illustrative model simulation of secondary tauopathy using a high-resolution structural brain connectome domain.

S8 Data: A low-resolution graph of the structural brain connectome. The graph is expressed in a standard format (graphml) based on the human-readable XML markup language. This graph consists of 83 vertices (anatomical regions of interest) and 1,654 edges; data for the graph was sourced from freely-available patient connectome data (https://braingraph.org).

S9 Data: A high-resolution graph of the structural brain connectome. The graph is expressed in a standard format (graphml) based on the human-readable XML markup language. This graph consists of 1,015 vertices (anatomical regions of interest) and 70,892 edges; data for the graph was sourced from freely-available patient connectome data (https://braingraph.org).

